# *PSEN1* Expression Identifies a Developmentally Distinct Favorable-Prognosis State in SHH α Medulloblastoma

**DOI:** 10.64898/2026.08.21.746295

**Authors:** Julia Vanini, Amanda Thomaz, Marina Müller Lupatini, André T. Brunetto, Caroline Brunetto de Farias, Mariane Jaeger, Rafael Roesler

## Abstract

**Background:** Although *PSEN1* is best known for its role in Alzheimer’s disease, it also regulates neural development and cerebellar morphogenesis. Medulloblastoma (MB) is the most common malignant pediatric brain tumor and arises from disrupted cerebellar developmental programs. The clinical significance of *PSEN1* in MB remains unknown. We investigated the prognostic value and transcriptional correlates of *PSEN1* expression across molecular subgroups and subtypes of MB.

**Methods:** Public bulk and single-cell transcriptomic datasets were used to examine *PSEN1* expression, associations with overall survival (OS), and transcriptional correlates in MB. The SHH α-associated transcriptional pattern was evaluated in an independent cohort, and *PSEN1* expression was further examined in the developing human cerebellum and across pediatric brain tumor types. Genes strongly correlated with *PSEN1* in SHH α MB were subjected to Gene Ontology (GO) enrichment analysis.

**Results:** High *PSEN1* expression was consistently associated with significantly longer OS exclusively in SHH α MB. The *PSEN1*-associated transcriptional pattern was reproduced in an independent SHH α cohort. *PSEN1* was expressed across developing cerebellar cell populations and pediatric brain tumor types, with MB showing intermediate expression among the tumor entities examined. In SHH α MB, *PSEN1* was associated with a coordinated transcriptional program enriched for RNA homeostasis, intracellular membrane trafficking, protein quality control, lipid and calcium signaling, and developmental pathways.

**Conclusions:** High *PSEN1* expression identifies a favorable-prognosis subset of SHH α MB and is associated with a distinct transcriptional program related to endomembrane organization and cellular homeostasis rather than canonical SHH signaling. These findings suggest that *PSEN1* may mark a developmentally distinct tumor state and generate new hypotheses regarding subtype-specific developmental programs in MB.

**Graphical Abstract:** 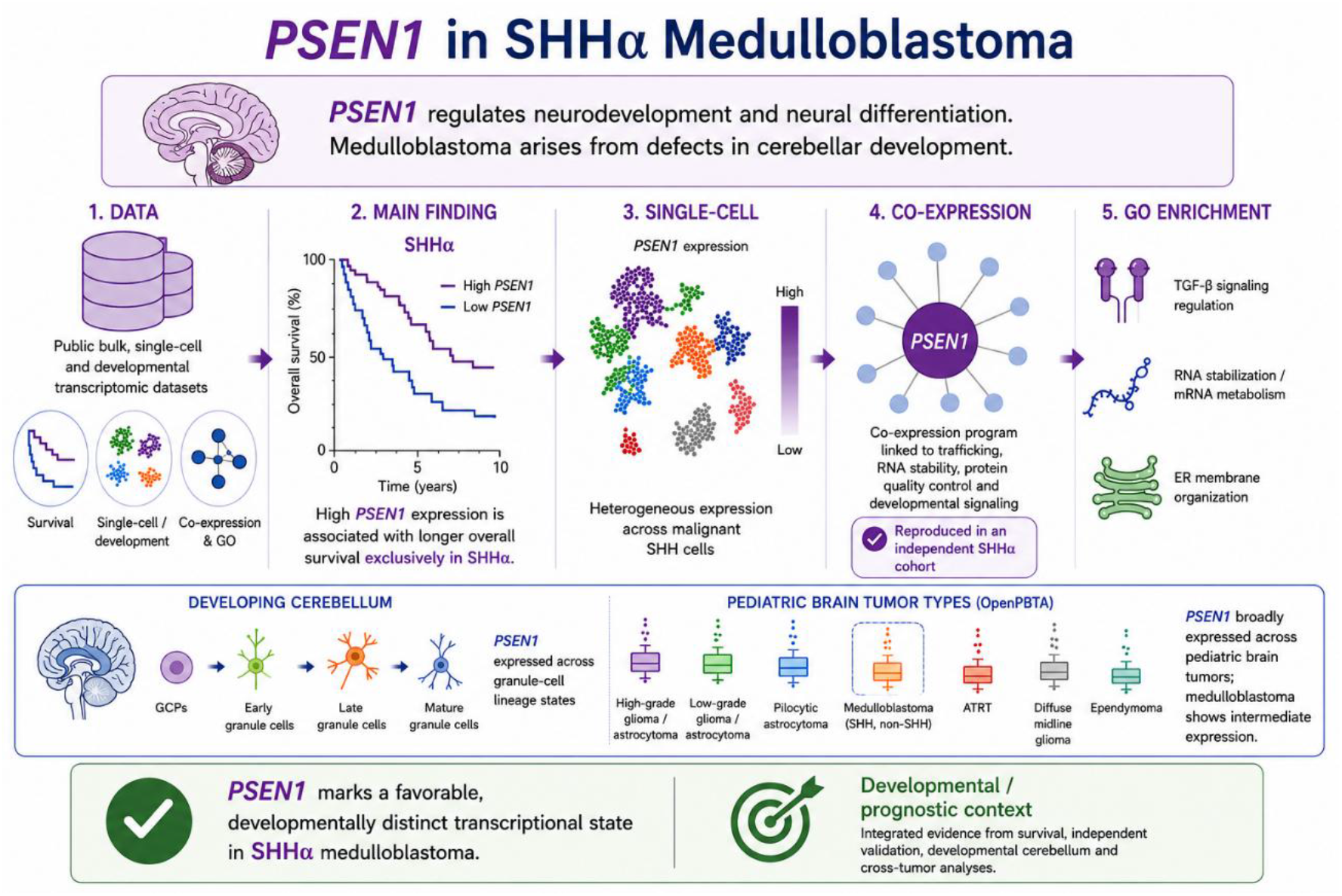

High *PSEN1* expression identifies a favorable-prognosis state in SHH α medulloblastoma and is associated with a reproducible transcriptional program related to cellular homeostasis and development. Its broad expression during cerebellar development and across pediatric brain tumors suggests that its prognostic significance reflects the specific molecular context of SHH α tumors.

## 1 Introduction

Pediatric brain tumors represent the most common solid malignancies in childhood and remain a leading cause of cancer-related death among children. Medulloblastoma (MB) is the most frequently diagnosed malignant brain tumor, arising in the cerebellum and serving as a paradigm of how disrupted neurodevelopmental processes can drive oncogenesis. Current treatment relies on a combination of surgical resection, radiotherapy, and chemotherapy, which has substantially improved survival rates over recent decades. Nevertheless, disease recurrence still occurs in a significant proportion of patients and is associated with a poor prognosis. Furthermore, treatment-related toxicities often result in persistent neurocognitive, neurological, and endocrine complications that can substantially affect long-term quality of life (Northcott et al. 2019).

Our understanding of the complexity of MB biology has consistently advanced thanks to genomic approaches. It is now established that MB comprises four biologically distinct entities that present different clinical outcomes. These defined molecular subgroups are Wingless (WNT)-activated, Sonic Hedgehog (SHH)-activated, Group 3, and Group 4 MB (Taylor et al. 2012; Northcott et al. 2017; Juraschka and Taylor 2019).

Given the substantial heterogeneity within each subgroup, they are currently further subdivided into subtypes. Through the Medulloblastoma Advanced Genomics International Consortium, Cavalli et al. (2017) assembled a cohort of 763 primary MB samples and applied similarity network fusion (SNF) and spectral clustering within each of the four subgroups across the cohort to integrate gene expression and DNA methylation profiling and generate a genome-wide profile. This led to the identification and characterization of twelve subtypes that show distinct somatic copy-number aberrations and clinical features, two of them within the WNT subgroup, four in SHH, three in Group 3, and three in Group 4 tumors.

The *PSEN1* gene encodes presenilin-1 (PS1), the catalytic subunit of the γ-secretase complex, a multi-protein protease responsible for cleaving numerous transmembrane substrates, notably amyloid precursor protein (APP). Through its role in APP processing, PS1 contributes to the generation of amyloid-β (Aβ) peptides. *PSEN1* is widely regarded as a major Alzheimer’s disease (AD) gene, as pathogenic *PSEN1* mutations represent the most common cause of autosomal dominant early-onset familial AD (Sherrington et al. 1995). However, it also regulates neural development and differentiation. Lack of *PSEN1* leads to early differentiation of neural progenitor cells, leading to abnormal neuronal migration and disorganization of the laminar architecture in the developing brain (Handler et al. 2000). *PSEN1* mutations associated with familial AD result in premature neurogenesis in brain organoids derived from human induced pluripotent stem cells (iPSCs) (Arber et al 2021). In the cerebellum, *PSEN1* inactivation disrupts the spread of granule cell precursors (GCPs) to form the external granule layer (EGL) during development (Louvi et al. 2004).

Previous studies have not examined a possible role for *PSEN1* or presenilin-1 in MB. In this study, we used the Cavalli cohort, first screening each MB subtype for possible associations between *PSEN1* expression levels and patient outcome indicated by overall survival (OS). Of all twelve subtypes, only SHH α had a consistent statistically significant association with better OS. Among the four SHH MB subtypes (SHH α, SHH β, SHH γ, and SHH δ), SHH α is biologically the most genomically unstable and clinically aggressive subtype. Afflicting mostly children aged 3–16 years, SHH α tumors are characterized by amplifications of *MYCN*, *GLI2*, and *YAP1*, loss of 9q, 10q, and 17p, and a high rate of *TP53* mutations that define one of the worst prognoses among MB subtypes (Northcott et al., 2012; Cavalli et all., 2017; Coltin et al. 2021; Skowron et al. 2021; Gold et al. 2024). We then aimed to explore the expression and co-expression structure of *PSEN1* in SHH α.

## 2 Methods

### 2.1 *PSEN1* Gene Expression Across MB Subgroups and Subtypes

The Pomeroy dataset established by Cho et al. (2011; accession number GSE202043) was used to compare *PSEN1* expression in MB tumors (*n* = 194) versus non-tumoral cerebellar tissue (CB, *n* – 11). *PSEN1* mRNA expression across MB molecular subgroups and subtypes was evaluated using the GlioVis data portal (https://gliovis.bioinfo.cnio.es/) (Bowman et al. 2017). Analyses were performed using the Cavalli MB dataset (Cavalli et al. 2017; accession number GSE85217), which comprises 763 primary MB samples classified into the four distinct molecular subgroups and twelve molecular subtypes, and the platform’s built-in plotting tools were used to visualize *PSEN1* expression distributions among molecular subgroups and subtype-specific cohorts. The comparison between MB tumors and CB was performed with a two-tailed Mann–Whitney U test. Comparisons among subgroups and subtypes were performed using Tukey’s Honest Significant Difference (HSD) tests.

### 2.2 *PSEN1* Expression in MB Single-Cell RNA Sequencing Data

*PSEN1* expression at the single-cell level was evaluated using MB single-cell RNA sequencing (scRNA-seq) data from the dataset GSE156053 (Riemondy et al. 2022), available through the Gene Expression Omnibus (GEO) repository. To investigate the distribution of *PSEN1* expression across MB cellular populations and SHH molecular subtypes, data were explored using the Pediatric Neuro-oncology Cell Atlas interactive platform (https://www.pneuroonccellatlas.org/). Uniform Manifold Approximation and Projection (UMAP) embeddings were used to visualize *PSEN1* expression patterns at single-cell resolution across the different MB subgroups.

### 2.3 Survival Analysis

Associations between gene expression and OS were evaluated using the R2 Genomics Analysis and Visualization Platform (https://hgserver1.amc.nl/). Kaplan–Meier survival analyses were performed using the platform’s default settings. Patients were stratified into low- and high-expression groups according to two approaches: (1) the median expression value and (2) the optimal expression cutoff determined by the R2 scan procedure, which systematically evaluates potential cutoff values and selects the threshold yielding the strongest separation between survival curves.

Statistical significance was assessed using the log-rank test. For analyses based on the optimal scan cutoff, both the raw log-rank *p*-values and the multiple-testing-adjusted *p*- values provided by the platform were recorded, including Bonferroni-corrected and Benjamini–Hochberg (BH) false discovery rate (FDR) adjusted *p*-values. Results obtained using the median cutoff and scan cutoff approaches were compared to evaluate the robustness and consistency of survival associations across different patient stratification strategies.

### 2.4 Clinical Annotation Heatmap

To explore the relationship between *PSEN1* expression and clinicopathological features within the SHH α MB subtype, a heatmap was generated using the R2 Genomics Analysis and Visualization Platform. Samples were ordered according to *PSEN1* expression levels, and corresponding clinical annotations were displayed alongside expression values. The clinical variables examined included age group, vital status (alive/deceased), metastatic status, disease status, and histological subtype. This analysis was performed for exploratory and visualization purposes to assess potential patterns linking *PSEN1* expression with clinical characteristics. No formal statistical testing was performed for the heatmap analysis, which was intended to provide a visual overview of the distribution of clinical features across *PSEN1* expression levels.

### 2.5 *PSEN1* Co-Expression Structure

Genes co-expressed with *PSEN1* were identified using Spearman correlation analysis in SHH α MB samples. Genes exhibiting strong positive correlations with *PSEN1* (*r* ≥ 0.62) were ranked, and the 26 most strongly correlated genes were selected for visualization. A strong correlation was defined as *r* between 0.62 and 0.88 with a statistically significant *p* value, and a very strong correlation was defined as *r* > 0.89 with a statistically significant *p* value. These thresholds and ranges were chosen to retain genes showing consistently strong positive associations with *PSEN1* while maintaining a sufficiently large gene set for meaningful functional enrichment analysis. Correlation coefficients in this range are commonly interpreted as representing strong positive relationships in biomedical research. A heatmap displaying the expression profiles of *PSEN1* and these genes across tumor samples was generated in R. Expression values were visualized using unsupervised hierarchical clustering of genes and samples to identify patterns of coordinated transcriptional activity associated with *PSEN1* expression.

### 2.6 Functional Enrichment Analysis

To investigate biological processes associated with *PSEN1* expression in SHH α MB, gene co-expression analysis was performed using the R2 Genomics Analysis and Visualization Platform in the Cavalli cohort. Analyses were restricted to the SHH α molecular subtype. Genes exhibiting strong to very strong positive correlations with *PSEN1* expression (Spearman correlation coefficient *r* ≥ 0.62) were selected, yielding a set of 26 strongly co-expressed genes. This threshold was selected based on commonly used interpretations of correlation strength in biomedical research, in which coefficients above approximately 0.60 are generally considered strong positive correlations and approach the range often described as very strong correlations. The threshold was chosen to prioritize biologically meaningful associations while maintaining adequate sensitivity for downstream analyses of co-expression networks.

Functional enrichment analysis of the resulting gene set was performed using Enrichr (https://maayanlab.cloud/Enrichr/). Gene Ontology enrichment was assessed across the Biological Process (BP), Cellular Component (CC), and Molecular Function (MF) categories. Enriched terms were ranked according to the significance metrics provided by the platform, and terms with adjusted *p*-values < 0.05 were considered statistically significant.

### 2.7 Independent Validation of *PSEN1* Expression and Co-Expression Patterns

To independently assess the molecular findings obtained in the Cavalli cohort, gene-expression data from an additional MB dataset were analyzed using the harmonized expression matrix reported by Arora et al. (2026) Molecular subtype annotations for SHH MB were obtained from the Skowron et al. cohort (2021). Sample identifiers were matched between datasets, yielding 196 SHH tumors with available molecular subtype classification, comprising 50 SHH α, 42 SHH β, 32 SHH γ, and 72 SHH δ tumors. *PSEN1* expression was compared across SHH molecular subtypes using the Kruskal–Wallis test, followed by pairwise Wilcoxon rank-sum tests with BH correction for multiple comparisons.

To independently evaluate the *PSEN1*-associated transcriptional pattern identified in the Cavalli SHH α cohort, analyses were restricted to the 50 independently classified SHH α tumors. Pearson correlation coefficients between *PSEN1* expression and genome-wide gene expression were calculated. Of the 26 genes previously identified as strongly positively correlated with *PSEN1* in Cavalli SHH α tumors, 22 were represented in the independent expression matrix and were therefore evaluated as a prespecified validation gene set. Enrichment of these genes among the 100 genes showing the strongest positive correlations with *PSEN1* was assessed using a hypergeometric test. In addition, an empirical permutation analysis was performed by randomly sampling 100,000 sets of 22 genes from the genome-wide background distribution and comparing their median *PSEN1* correlation coefficients with that of the prespecified validation set. All analyses were performed in R (version 4.5.3) using RStudio (version 2026.08.1+195; Posit Software, PBC).

### 2.8 Developing Human Cerebellum Single-Nucleus Transcriptomic Analyses

*PSEN1* expression during normal human cerebellar development was examined in two independent single-nucleus RNA-sequencing datasets. First, we analyzed the developing human cerebellum atlas reported by Aldinger et al. (2021), comprising 69,174 nuclei sampled across 9–21 post-fertilization weeks. Cell-type and developmental-stage annotations provided with the dataset were retained. *PSEN1* expression was summarized across annotated cell populations as mean expression and the percentage of nuclei with detectable expression. Expression was also examined across developmental stages.

To specifically assess the granule-cell developmental lineage, nuclei annotated as rhombic lip (RL), GCP, and granule neuron (GN) were analyzed. Because nuclei obtained from the same donor are not independent biological replicates, statistical comparisons among these lineage states were performed using donor-level mean *PSEN1* expression. Donors represented in all three states were included in the paired analysis, and differences among RL, GCP, and GN were assessed using the Friedman test.

As an independent validation, *PSEN1* expression was additionally examined in the developing human cerebellum single-nucleus transcriptomic atlas reported by Sepp et al. (2024). Analyses were restricted to cells assigned by the authors to the RL/EGL developmental lineage. *PSEN1* expression was summarized across the author-defined developmental states as mean expression and percentage of nuclei with detectable expression. Processed single-nucleus datasets and accompanying cell annotations were obtained through CELLxGENE (https://cellxgene.cziscience.com/). All analyses were performed in R (version 4.5.3) using RStudio (version 2026.08.1+195; Posit Software, PBC).

### 2.9 *PSEN1* Expression Across Pediatric Brain Tumor Types

*PSEN1* expression across pediatric brain tumor types was examined using RNA-sequencing data from the Open Pediatric Brain Tumor Atlas (OpenPBTA; https://github.com/AlexsLemonade/OpenPBTA-analysis; Shapiro et al. 2023). Gene-level expression data were obtained from the stranded RNA-sequencing RSEM transcript-per-million (TPM) matrix from OpenPBTA release v23 (release-v23- 20230115) and matched to harmonized tumor annotations using Kids First Biospecimen identifiers. The analysis included seven pediatric brain tumor groups represented by at least 30 samples in the expression dataset: high-grade glioma/astrocytoma (*n* = 93), low-grade glioma/astrocytoma (*n* = 86), pilocytic astrocytoma (*n* = 131), MB (*n* = 121), atypical teratoid/rhabdoid tumor (ATRT; *n* = 30), diffuse midline glioma (*n* = 34), and ependymoma (EPN, *n* = 93). Differences in *PSEN1* expression across tumor groups were assessed using the Kruskal–Wallis test. Pairwise comparisons were subsequently performed using Wilcoxon rank-sum tests with BH correction for multiple comparisons. All analyses were performed in R (version 4.5.3) using RStudio (version 2026.08.1+195; Posit Software, PBC).

## 3 Results

### 3.1 High *PSEN1* Expression is Selectively Associated with Better Survival in SHH α MB

We first screened *PSEN1* expression separately in all twelve MB subtypes to determine associations with patient survival, using four complementary statistical approaches to determine whether associations were consistent across different methods of patient stratification. Across four different statistical criteria for significance, namely showing the raw *p*-value for analyses using the median as cutoff for separating low versus high expression; the Bonferroni-adjusted *p*-value with the R2 scan tool cutoff; the raw *p*-value with the scan cutoff; and FDR-adjusted *p*-value obtained using the BH method to adjust for multiple comparisons, with the scan cutoff, the only subtype showing significant associations between *PSEN1* expression and survival in all statistical methods was SHH α, where *PSEN1* transcription showed a consistent correlation with longer OS (Figure 1, Supplementary Table S1).

**FIGURE 1.**
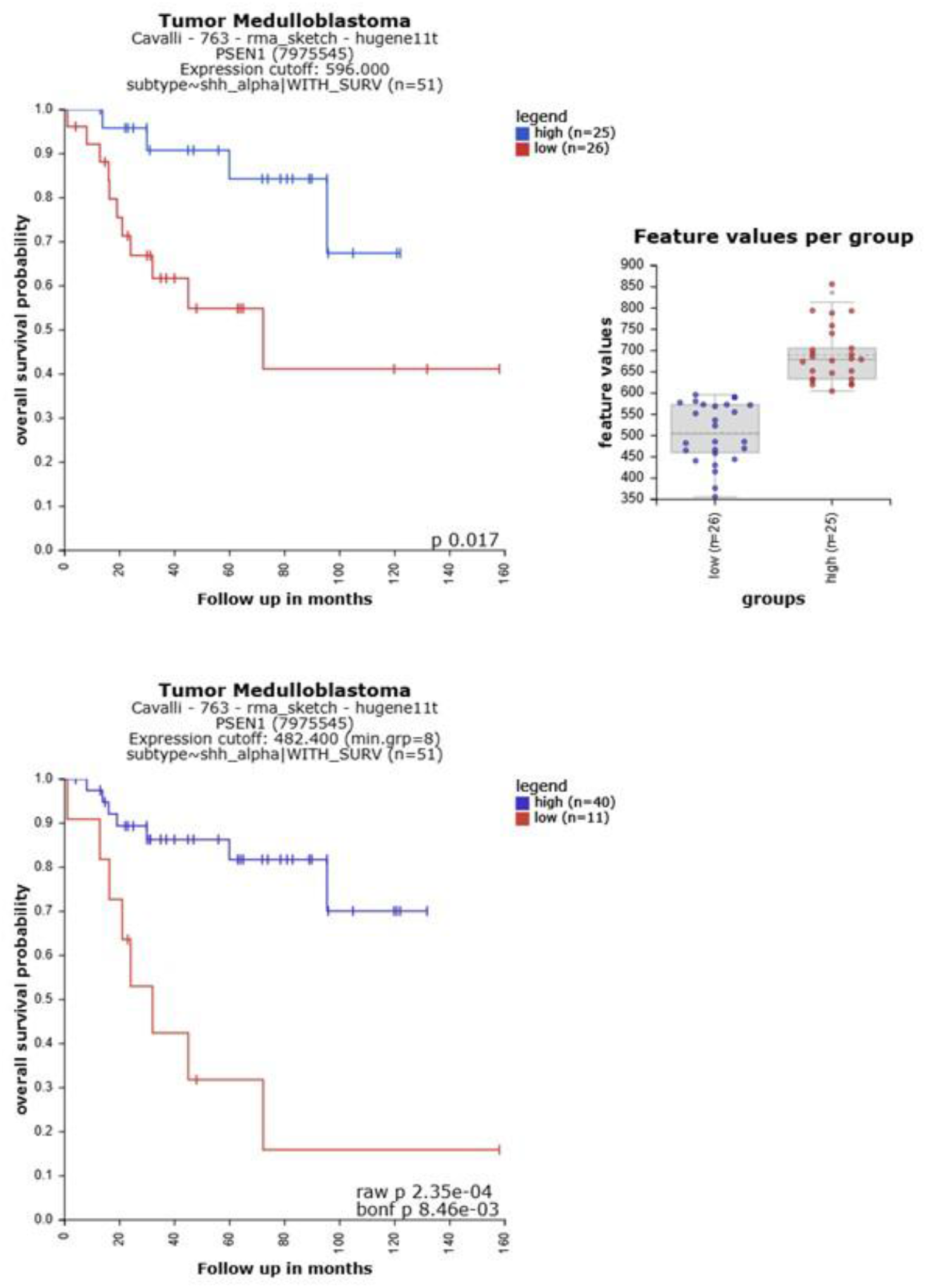
Kaplan-Meier analyses of OS according to *PSEN1* gene expression levels in SHH α MB tumors. Survival analyses were performed using the R2 Genomics Analysis and Visualization Platform. Upper panels, results using the raw *p* value with median cutoff. Lower panel, raw and Bonferroni-adjusted *p* values using optimal scan cutoff; *p* values are shown in the panels and Supplementary Table S1.

### 3.2 *PSEN1* Expression Levels in MB Tumors Classified According to Molecular Subgroup and Subtype

*PSEN1* transcription levels were significantly higher in MB tumors in comparison with non-tumoral CB (Figure 2). Among molecular subgroups, SHH tumors exhibited significantly lower *PSEN1* expression than WNT tumors, whereas WNT tumors showed significantly higher expression than Group 4 tumors (Figure 3, Supplementary Table S2). No other subgroup comparisons reached statistical significance. At the subtype level, SHH α tumors displayed significantly lower *PSEN1* expression than WNT-α and Group 3-α tumors (Figure 3, Supplementary Table S3). No significant differences were observed between SHH α and the remaining subtypes.

**FIGURE 2.**
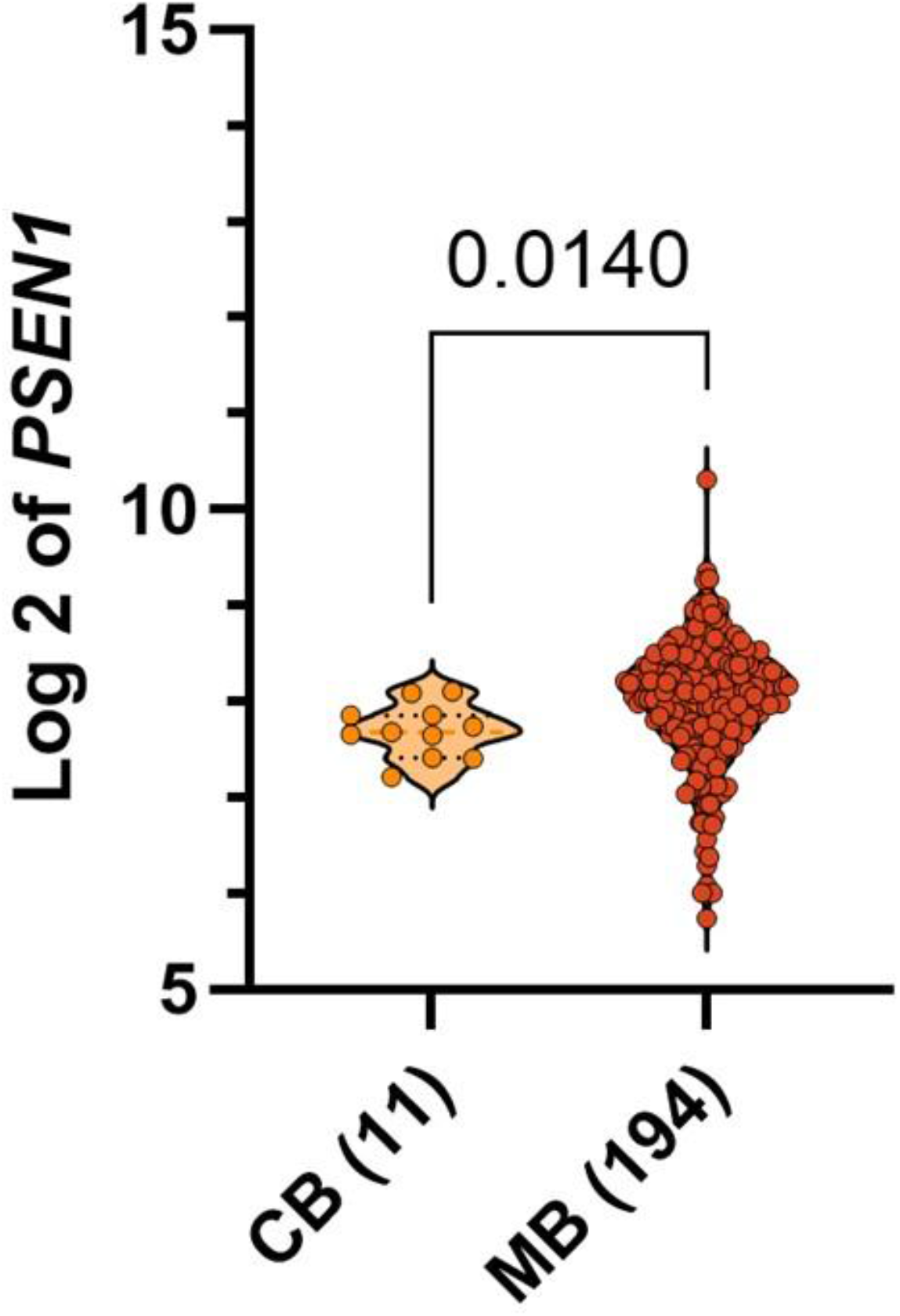
*PSEN1* expression in MB tumors and non-tumoral cerebellar tissue (CB). Transcription levels were compared between CB (*n* = 11) and MB samples (*n* = 194). Data were obtained from the Pomeroy dataset (GSE202043) described by Cho et al. (2011). Each point represents an individual sample. Statistical significance was assessed using a two-tailed Mann–Whitney U test, with the *p* value indicated above the comparison. The bracket denotes the statistically significant difference between groups. Data are presented as log₂-transformed *PSEN1* mRNA level values.

**FIGURE 3.**
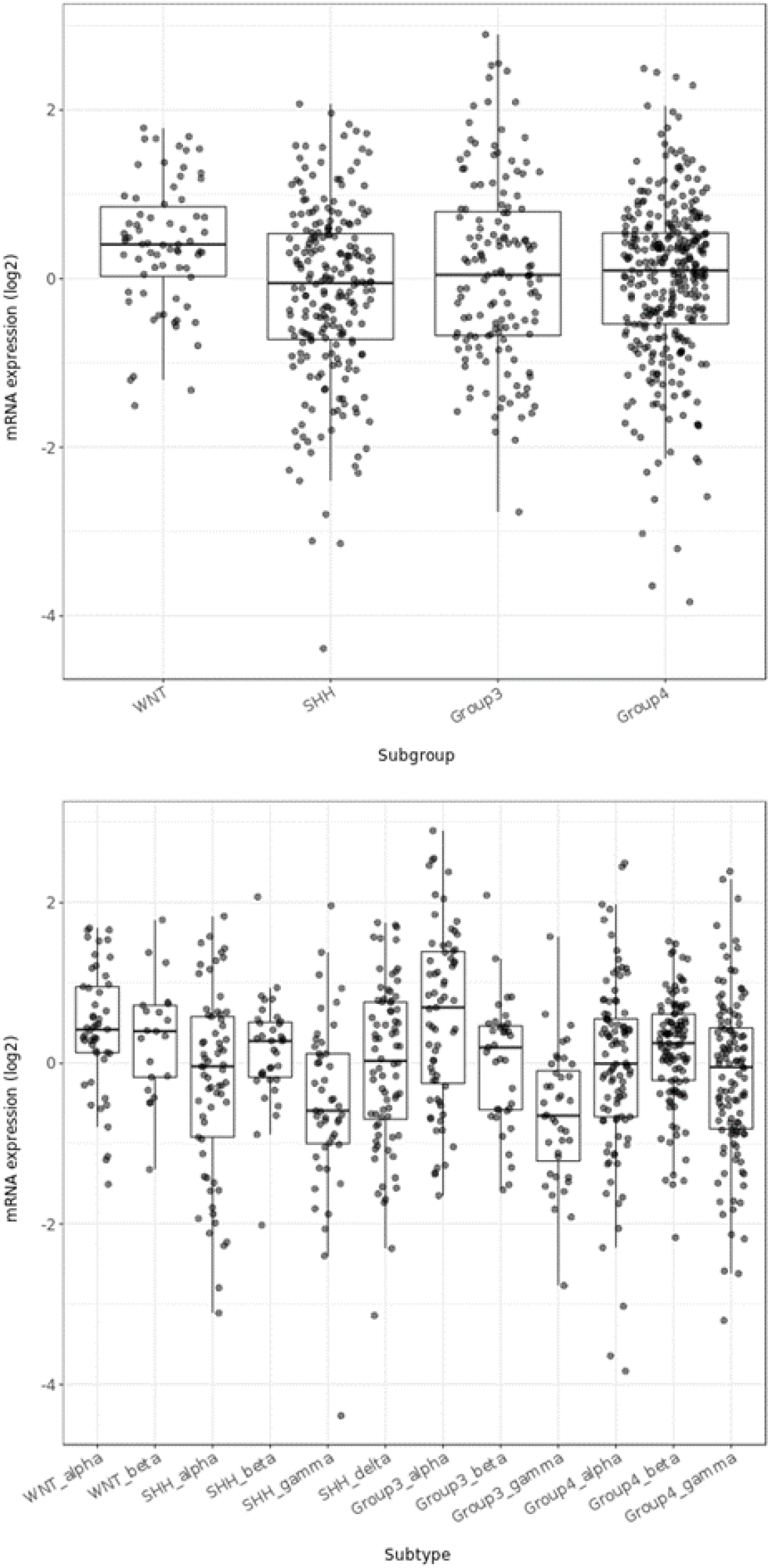
*PSEN1* expression across MB molecular subgroups and subtypes in the Cavalli cohort. Upper panel, comparison among the four molecular subgroups. Lower panel, comparison among the twelve molecular subtypes. Statistical analyses were performed using the Tukey’s Honest Significant Difference (HSD) test and *p* values are shown in Supplementary Tables S2 and S3.

To investigate *PSEN1* expression at the single-cell level, scRNA-seq data from SHH MB samples were analyzed. UMAP visualization demonstrated heterogeneous *PSEN1* expression across malignant cell populations, with *PSEN1*-positive cells distributed throughout the SHH tumor cell landscape rather than being restricted to a discrete cellular cluster (Figure 4). Within the SHH α samples from the Cavalli cohort (*n* = 65), *PSEN1* expression scores were further visualized according to age group, survival status, histological subtype, and metastatic status (Supplementary Figure S1).

**FIGURE 4.**
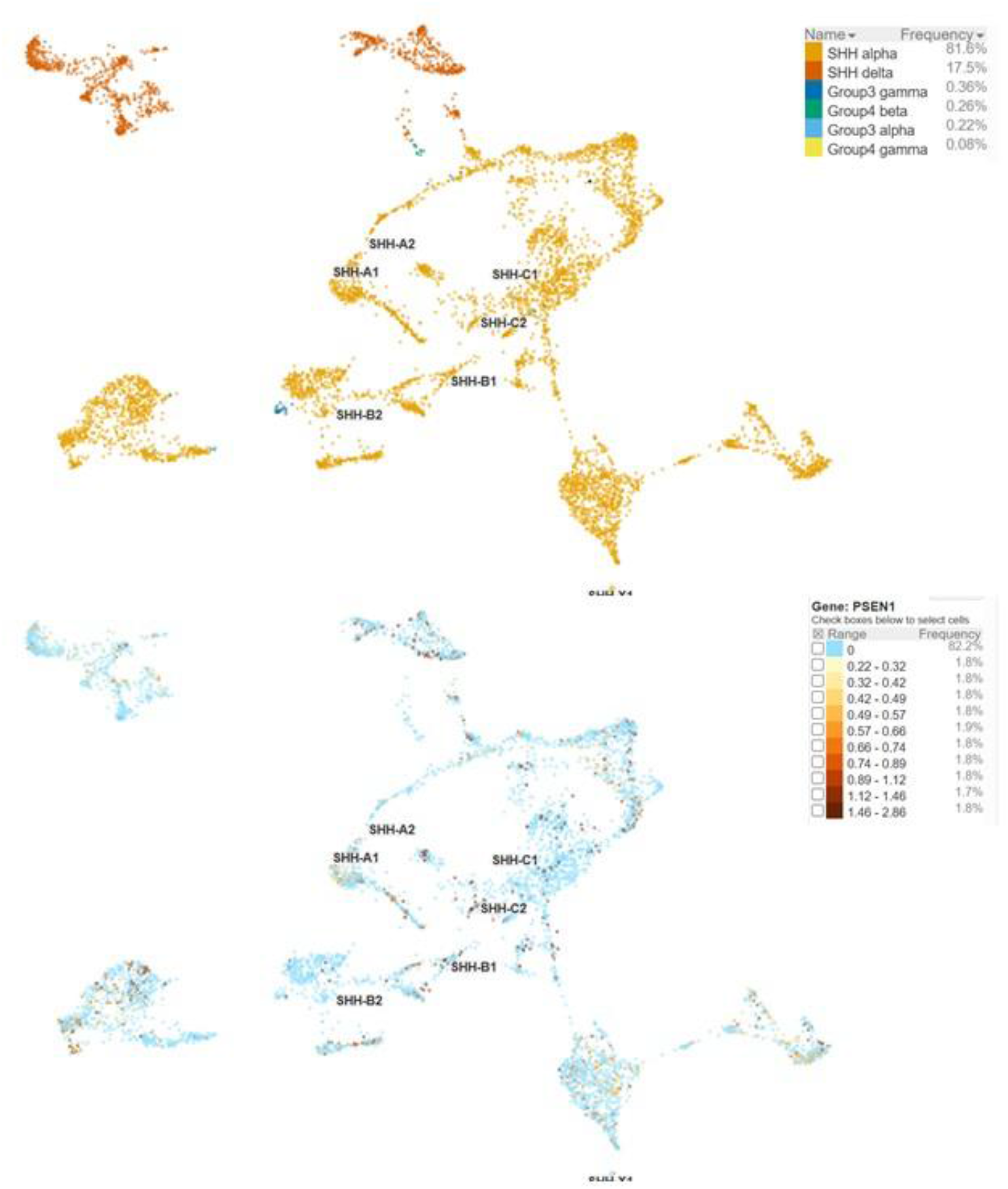
UMAP visualization of *PSEN1* expression in single-cell RNA sequencing data from SHH MB samples. *PSEN1*-positive cells are distributed throughout malignant SHH tumor populations and subtypes rather than being confined to a discrete cellular cluster. Labels refer to single-cell transcriptional states identified by clustering individual malignant cells across SHH tumors (Riemondy et al. 2022).

### 3.3. *PSEN1* Co-Expression Network and Functional Enrichment Analysis in SHH α MB

To explore candidate biological processes associated with *PSEN1* expression in SHH α MB, co-expression analyses were performed using transcriptomic data from the Cavalli cohort. Twenty-six genes exhibited strong or very strong positive correlations with *PSEN1* expression and were selected for subsequent analyses. The strongest correlations were observed for *SPTLC2*, *ATL1*, *EXD2*, *ZNF410*, *PAPOLA*, *SOS2*, *EXOC5*, *VIPAR*, *SEL1L*, *SLC39A9*, *MUDENG*, *C14ORF118*, *TMED8*, *SETD3*, *WDR20*, *C14ORF101*, *SNX6*, *DCAF5*, *GOLGA5*, *PPM1A*, *ACTR10*, *TRAPPC6B*, *SNW1*, *ZFYVE1*, *ZC3H14*, and *JKAMP* (Supplementary Table S4). Hierarchical clustering demonstrated a coordinated expression pattern among *PSEN1* and its co-expressed genes across SHH α tumors (Figure 5).

**FIGURE 5.**
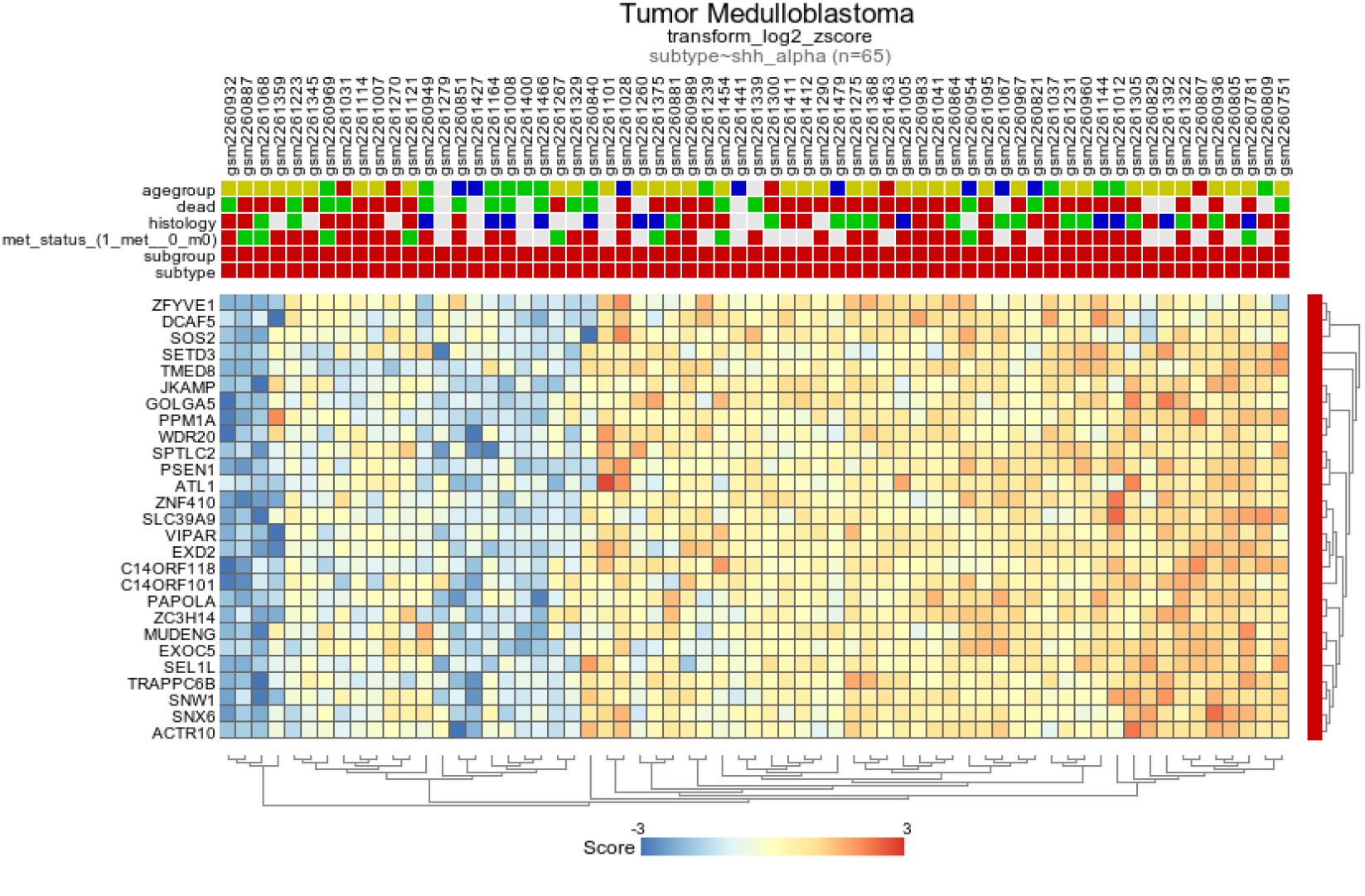
Heatmap showing coordinated expression of *PSEN1* and the 26 top genes displaying the strongest positive correlations with *PSEN1* in SHH α MB. Expression values are represented as log2-transformed z-scores. Both genes and tumor samples were hierarchically clustered using unsupervised clustering.

Gene Ontology enrichment analysis of the *PSEN1* co-expression signature identified significant enrichment of terms related to RNA homeostasis, protein processing, intracellular trafficking, and membrane-associated cellular compartments. Within the Biological Process category, the most significantly enriched terms included regulation of transforming growth factor β receptor signaling pathway, RNA stabilization, mRNA stabilization, negative regulation of mRNA catabolic process, and regulation of mRNA stability. Cellular Component analysis identified enrichment of structures associated with intracellular membrane trafficking and protein quality control, including the Hrd1 ubiquitin ligase ERAD-L complex. Molecular Function analysis revealed enrichment of activities related to sphingolipid biosynthesis, calcium/calmodulin-dependent phosphatase signaling, ubiquitin-mediated protein quality control, RNA processing, DNA repair, and epigenetic regulation. These findings suggest that the *PSEN1*-associated transcriptional program is linked to neuronal membrane homeostasis, intracellular calcium signaling, and maintenance of differentiated cellular functions rather than to proliferative pathways (Figure 6).

**FIGURE 6.**
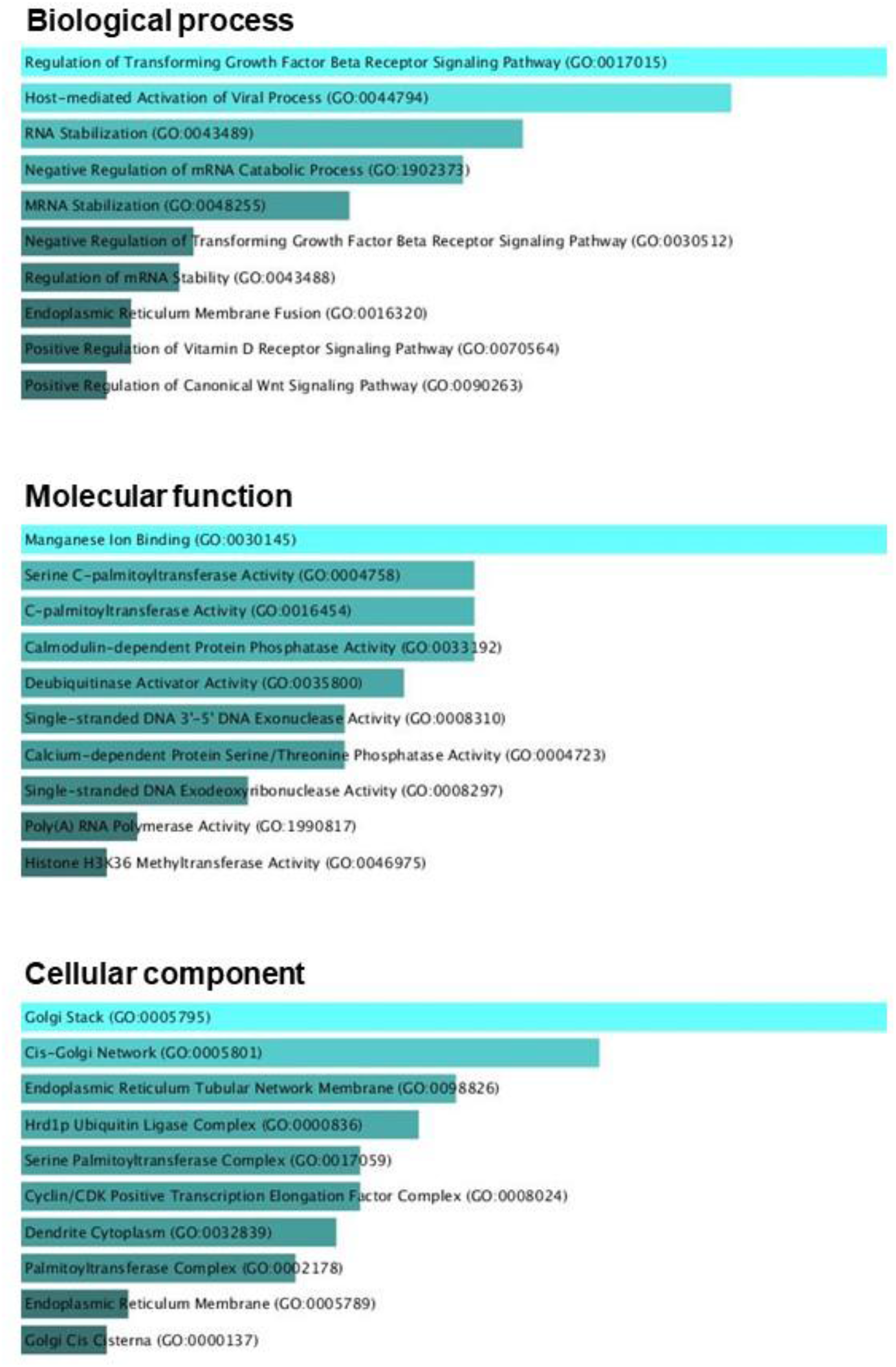
Gene Ontology enrichment analysis of the 26 genes most strongly correlated with *PSEN1* in SHH α MB. Biological Process (BP), Cellular Component (CC), and Molecular Function (MF) analyses demonstrate enrichment for RNA homeostasis, intracellular membrane trafficking, protein quality control, developmental signaling pathways, and neuronal membrane-associated functions.

Together, these findings demonstrate that elevated *PSEN1* expression in SHH α MB is associated with a transcriptional program enriched for genes involved in RNA regulation, intracellular membrane systems, protein quality control, and developmental signaling pathways.

### 3.4 Independent Validation of *PSEN1*-Associated Molecular Features in SHH MB

We next examined *PSEN1* expression and its associated transcriptional pattern in an independent cohort of molecularly characterized SHH MB (Arora et al. 2026; Skowron et al. 2021). A total of 196 tumors could be assigned to SHH molecular subtypes, including 50 SHH α, 42 SHH β, 32 SHH γ, and 72 SHH δ tumors. *PSEN1* expression differed modestly across the four SHH subtypes (*p* = 0.041). Median expression values were 3.220, 3.405, 2.960, and 3.115 in SHH α, SHH β, SHH γ, and SHH δ, respectively. In pairwise analyses, only the difference between SHH β and SHH γ remained significant after BH correction (adjusted *p* = 0.026), whereas SHH α did not differ significantly from the other individual subtypes (Supplementary Figure S2). Thus, consistent with the Cavalli dataset analysis, the distinctive association of *PSEN1* with SHH α biology does not appear to result simply from selectively elevated *PSEN1* expression in this subtype.

We then asked whether the *PSEN1*-associated transcriptional pattern identified in Cavalli SHH α tumors could be reproduced in the independent set of 50 SHH α tumors. Of the 26 genes comprising the Cavalli-derived *PSEN1*-associated gene set, 22 were represented in the independent expression matrix. Remarkably, all 22 genes showed positive correlations with *PSEN1*, with Pearson correlation coefficients ranging from 0.674 to 0.944 and a median correlation of 0.880 (Supplementary Figure S3). Fifteen of the 22 genes ranked among the 100 genes most strongly correlated with *PSEN1* genome-wide, representing significant enrichment relative to chance (hypergeometric *p* = 1.29 × 10⁻²⁹). Moreover, the median correlation of the prespecified 22-gene set exceeded that obtained for all 100,000 randomly sampled gene sets of equal size (empirical *p* < 1 × 10⁻⁵). These findings independently support the reproducibility of the *PSEN1*-associated transcriptional pattern observed in SHH α MB.

### 3.5 *PSEN1* is Expressed Across Cell Populations of the Developing Human Cerebellum

To characterize the developmental context of *PSEN1* expression, we first analyzed a single-nucleus transcriptomic atlas comprising 69,174 nuclei from the developing human cerebellum across 9–21 post-fertilization weeks (Aldinger et al. 2021). *PSEN1* was detected across multiple cerebellar cell populations, with the highest mean expression observed in endothelial cells, brainstem choroid/ependymal cells, and pericytes (Figure 7A), indicating that its expression is not restricted to a specific neuronal lineage. Importantly, *PSEN1* was detected throughout the RL–granule cell lineage, including RL cells (mean expression, 0.211; 15.2% *PSEN1*-positive nuclei), GCPs (0.217; 14.0%), and GNs (0.133; 10.1%). *PSEN1* expression was detectable throughout the sampled prenatal developmental period, with descriptively lower mean expression at several later developmental stages (Figure 7B). Because developmental stage is associated with differences in donor and cellular composition, this pattern was considered descriptive rather than evidence of longitudinal developmental regulation.

**FIGURE 7.**
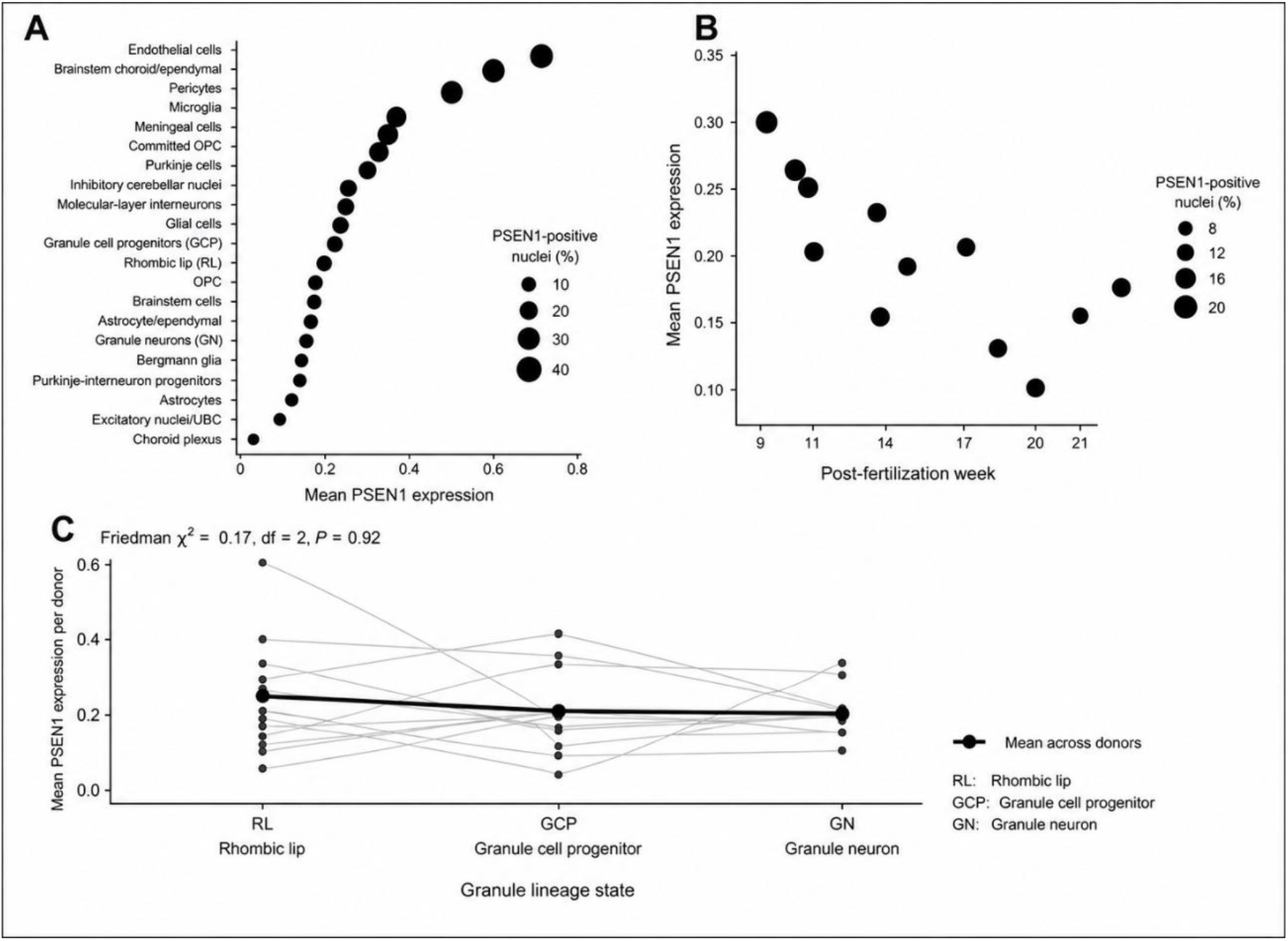
*PSEN1* expression in the developing human cerebellum. Single- nucleus RNA-sequencing data from the developing human cerebellum were analyzed using the dataset of Aldinger et al. (2021). **(A)** *PSEN1* expression across 21 annotated cerebellar cell populations. Dot position indicates mean *PSEN1* expression and dot size indicates the percentage of nuclei with detectable *PSEN1* expression. **(B)** Mean *PSEN1* expression across post-fertilization developmental stages. Dot size indicates the percentage of *PSEN1*-positive nuclei. Developmental-stage values are presented descriptively because developmental age is associated with differences in donor and cellular composition. **(C)** Donor-level mean *PSEN1* expression across the RL–GCP–GN developmental lineage. Thin lines connect values from the same donor, whereas the thick line and large points indicate mean expression across donors. No significant difference was observed among lineage states (Friedman χ² = 0.17, df = 2, *p* = 0.92).

We next compared *PSEN1* expression across the RL–GCP–GN lineage using donor-level mean expression to account for the non-independence of nuclei from the same individual. No significant difference was observed among the three lineage states (Friedman χ² = 0.17, df = 2, *p* = 0.92; Figure 7C), indicating that *PSEN1* is expressed across this developmental trajectory without evidence of preferential expression at a particular granule-lineage state.

We then examined an independent developing human cerebellum single-nucleus dataset from Sepp et al. (2024). *PSEN1* expression was again detected across author-defined RL/EGL developmental populations (Supplementary Figure S4). In particular, *PSEN1* was readily detected in GCPs (*n* = 2,714; mean expression = 0.237; 18.4% positive nuclei), early differentiating granule cells (GC_diff_1; *n* = 4,251; mean = 0.231; 18.3% positive), and defined granule cells (GC defined; *n* = 47,116; mean = 0.222; 17.4% positive). Expression was also detected in the GC_diff_2 state (*n* = 14,715; mean = 0.175; 14.8% positive). Thus, an independent developmental cerebellum dataset confirmed *PSEN1* expression across granule-cell progenitor and differentiating granule-cell states.

### 3.6 *PSEN1* is Expressed Across Different Pediatric Brain Tumor Types

To place *PSEN1* expression in MB within the broader context of pediatric brain tumors, we examined *PSEN1* transcript levels across seven tumor groups in the OpenPBTA dataset (Shapiro et al. 2023). The results are shown in Figure 8. *PSEN1* was expressed across all tumor types examined, but its expression differed significantly among tumor groups (Kruskal–Wallis χ² = 87.74, df = 6, *p* < 0.0001). Median *PSEN1* expression was 13.0 TPM in high-grade glioma/astrocytoma, 12.9 TPM in low-grade glioma/astrocytoma, 12.6 TPM in pilocytic astrocytoma, 11.1 TPM in MB, 10.9 TPM in ATRT, 10.6 TPM in diffuse midline glioma, and 9.21 TPM in EPN. Pairwise Wilcoxon rank-sum tests with BH correction showed that *PSEN1* expression in MB was significantly lower than in high-grade glioma/astrocytoma (adjusted *p* = 0.0069), low-grade glioma/astrocytoma (adjusted *p* = 9.12 × 10⁻⁵), and pilocytic astrocytoma (adjusted *p* = 4.70 × 10⁻⁴), and significantly higher than in EPN (adjusted *p* = 2.15 × 10⁻⁶). No significant differences were observed between MB and either ATRT or diffuse midline glioma (adjusted *p* = 0.434 for both comparisons). Thus, *PSEN1* is broadly expressed across pediatric brain tumors, with MB displaying intermediate expression levels among the tumor types examined.

**FIGURE 8.**
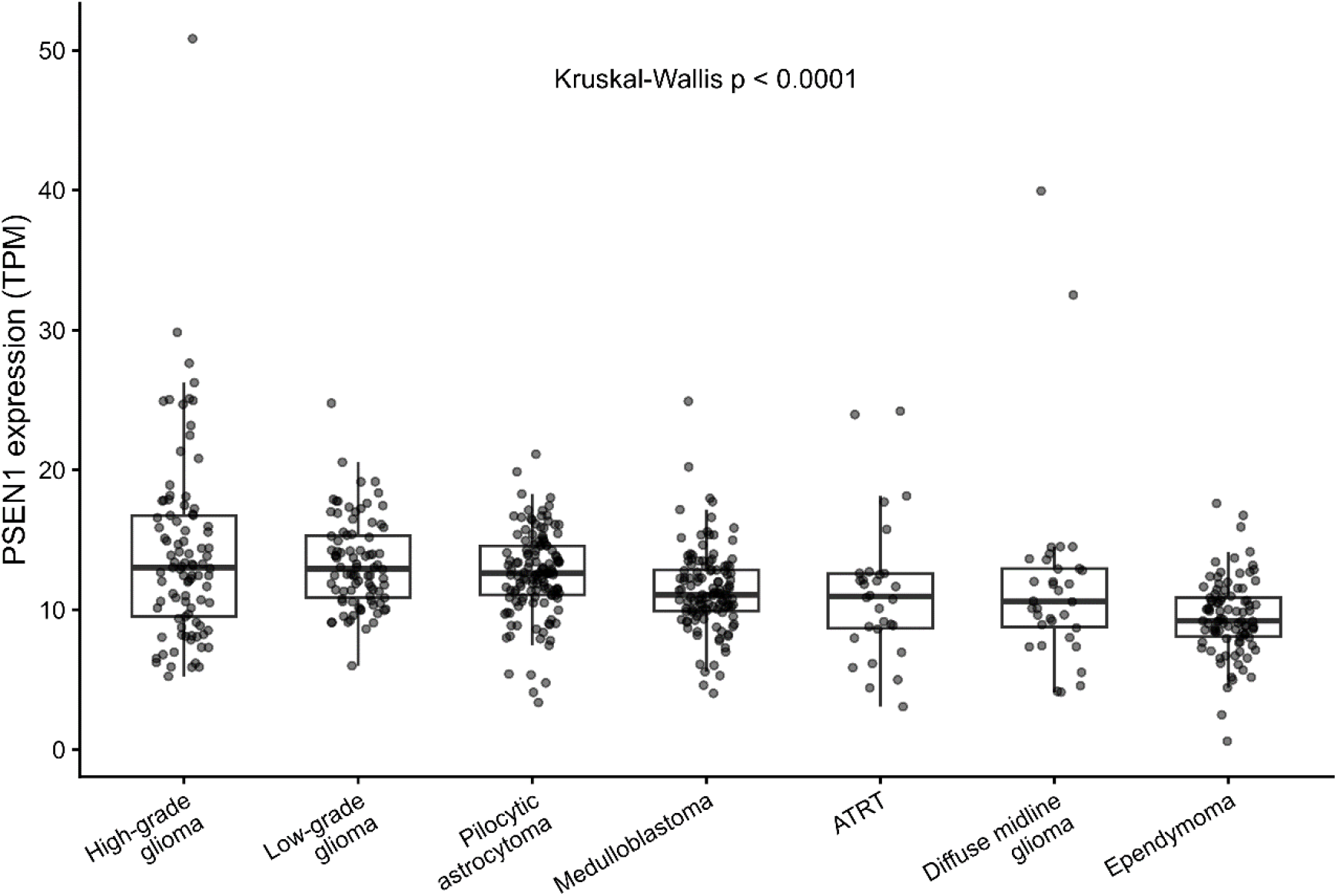
*PSEN1* expression across pediatric brain tumor types. *PSEN1* expression was examined using RNA-sequencing data from the OpenPBTA, including high-grade glioma/astrocytoma (*n* = 93), low-grade glioma/astrocytoma (*n* = 86), pilocytic astrocytoma (*n* = 131), MB (*n* = 121), ATRT (*n* = 30), diffuse midline glioma (n = 34), and EPN (*n* = 93). Expression levels are shown as TPM. Boxes represent the interquartile range, with the horizontal line indicating the median; individual points represent tumor samples. Differences among tumor types were assessed using the Kruskal–Wallis test (*p* < 0.0001). Pairwise comparisons were performed using Wilcoxon rank-sum tests with BH correction for multiple comparisons, and the results for these comparisons are described in the main text.

## 4 Discussion

Characterizing biological and clinical differences among MB subtypes is critical for improving risk stratification. SHH MB is thought to arise predominantly from GCPs located in the EGL of the developing cerebellum. These cells are derived from the upper RL (uRL) and are committed to a glutamatergic excitatory neuronal lineage. Tumorigenesis is typically initiated by activating alterations in the SHH signaling pathway, including mutations in *PTCH1*, *SMO*, and *SUFU*, whereas *MYCN* amplification is associated with a more aggressive clinical course and poorer outcome (Oliver et al. 2005; Yang et al. 2008; Gibson et al. 2010; Vladoiu et al. 2019; Gold et al. 2024; Falcón et al. 2026). Within the SHH subgroup, SHH α tumors are distinguished by marked chromosomal instability, extensive copy-number alterations, recurrent whole-chromosome arm gains and losses, frequent focal amplifications, and an unfavorable prognosis (Cavalli et al. 2017).

We have previously shown that several genes involved in neurodevelopment and neural plasticity can display different and often opposite patterns of association with patient survival in different molecular subgroups of MB (de Araújo et al. 2023; Fratini et al. 2023; Monteiro et al. 2024; Coelho et al. 2025; Saciloto et al. 2026). The present study highlights the importance of further investigating molecular targets and candidate prognostic markers at the subtype level within each molecular subgroup.

The proliferation of GCPs in the developing cerebellum is regulated by SHH signaling (Wallace 1999; Wechsler-Reya and Scott 1999). PS1 can enhance SHH expression and functionally interact with SHH to impair neurogenesis and promote cell proliferation and apoptosis (Paganelli et al., 2001). This evidence supports the possibility that PS1 is a direct modulator of SHH signaling, which could result in influences on SHH α MB aggressiveness. In neuronal progenitors and human iPSCs, mutations in *PSEN1* result in a premature differentiation phenotype, through a mechanism likely involving Notch signaling inhibition (Handler et al. 2000; Arber et al. 2021), suggesting that *PSEN1* contributes to the preservation of immature neural states. At first glance, our finding that elevated *PSEN1* expression is associated with improved survival appears counterintuitive, as stem cell-associated programs are often linked to tumor aggressiveness. However, single-cell studies have shown that MB contains cells spanning multiple developmental states, and the most aggressive populations are not always the least differentiated (Hovestadt et al. 2019; Riemondy et al. 2022). Therefore, the relationship between differentiation status and clinical behavior is more complex than a simple stemness- versus-differentiation paradigm. One possible explanation is that increased *PSEN1* expression within the SHH α MB subtype identifies tumors that remain developmentally constrained and retain transcriptional features of cerebellar progenitor populations, whereas reduced *PSEN1* expression may permit transition toward alternative neuronal differentiation programs associated with greater cellular plasticity and tumor fitness. In this model, a favorable prognostic association of *PSEN1* would not necessarily reflect a protective effect of stemness itself, but rather the preservation of a lineage-restricted developmental state. This hypothesis is consistent with emerging evidence indicating that MB progression is influenced not only by proliferative capacity but also by the acquisition of distinct developmental and neuronal programs (Ghasemi et al. 2024). Further studies integrating transcriptomic, proteomic, and single-cell analyses will be necessary to determine whether *PSEN1* actively regulates these cellular states or functions primarily as a marker of favorable tumor differentiation trajectories.

Importantly, *PSEN1* was not selectively or particularly highly expressed in SHH α tumors compared with other MB subtypes. Thus, the present findings should not be interpreted as indicating that *PSEN1* is an expression marker of SHH α MB. Rather, they suggest that the clinical significance of variation in *PSEN1* expression is context-dependent, with higher *PSEN1* expression being associated with favorable outcome specifically within the SHH α molecular background. Gene expression levels and the biological or clinical significance of their variation across individual tumors represent distinct phenomena; a gene need not be preferentially expressed in a tumor subtype for its expression to identify biologically or clinically distinct states within that subtype. The transcriptional programs associated with *PSEN1* in SHH α tumors further support the possibility that its prognostic association reflects the particular molecular context of these tumors. Nevertheless, the present data establish an association and do not demonstrate an SHH α-specific functional role for *PSEN1*, which will require validation in experimental models that faithfully reproduce this molecular subtype.

Our co-expression and GO analyses revealed an unexpected pattern of *PSEN1* co-expression in SHH α. The analyses did not recover a canonical SHH program or a classical neurodevelopmental program. Instead, most of the enriched terms converge on themes including endoplasmic reticulum, Golgi, membrane trafficking, proteostasis, and RNA homeostasis. Specific signaling pathways among the enriched terms included regulation of the transforming growth factor-β (TGF-β) receptor pathway and the canonical WNT pathway. The BP enrichment indicated RNA stabilization, mRNA stabilization, regulation of mRNA stability, and negative regulation of mRNA catabolism, whereas the CC analysis identified Hrd1 ubiquitin ligase complex. The findings may be interpreted as suggesting a cellular state dedicated to maintaining protein and transcript integrity. In other words, *PSEN1*-high SHH α MB may exhibit enhanced capacity for transcript maintenance, protein quality control, endoplasmic reticulum (ER)- associated degradation (ERAD), and trafficking of newly synthesized proteins. Regarding the developmental signaling pathways, TGF-β and WNT have known developmental interactions with *PSEN1* biology (Sanadgol et al. 2023; Lee et al. 2024). Overall, the results raise the possibility that high *PSEN1* expression in SHH α MB is associated with a cellular program featuring intracellular membrane trafficking, ER-Golgi function, protein quality control, and modulation of developmental signaling pathways, rather than with canonical SHH pathway activation itself.

The independent-cohort analysis further supports the association of *PSEN1* with a reproducible transcriptional state in SHH α MB. *PSEN1* expression was not selectively increased in SHH α compared with the other SHH molecular subtypes, supporting the interpretation that its association with favorable outcome in SHH α is unlikely to reflect simply greater expression of *PSEN1* in this subtype. More importantly, the *PSEN1*- associated gene set identified in Cavalli SHH α tumors showed striking concordance in an independent set of 50 SHH α tumors, so that all 22 evaluable genes were positively correlated with *PSEN1*, and 15 ranked among the 100 strongest genome-wide *PSEN1* associations. This enrichment remained highly significant when evaluated against the genome-wide correlation background and by random gene-set permutation. These findings suggest that the association of *PSEN1* with SHH α MB reflects a reproducible transcriptional context rather than a cohort-specific co-expression pattern.

Analysis of two independent single-nucleus transcriptomic datasets of the developing human cerebellum provided additional developmental context for the association between *PSEN1* expression and favorable outcome in SHH α MB. In the Aldinger et al. dataset, *PSEN1* was broadly expressed across cerebellar cell populations and was therefore not selectively associated with the granule lineage. Nevertheless, expression was clearly detectable in RL cells, GCPs, and GNs. Donor-level analysis did not demonstrate significant differences in *PSEN1* expression across the RL–GCP–GN trajectory, arguing against a simple model in which *PSEN1* marks a particular stage of granule-cell differentiation.

Analysis of an independent developing human cerebellum dataset from Sepp et al. (2024) confirmed *PSEN1* expression in GCPs and across differentiating granule-cell states. Together, these observations indicate that *PSEN1* is physiologically expressed throughout the developmental lineage relevant to SHH MB biology, although it is neither restricted to this lineage nor selectively enriched at a specific developmental state. The developmental data therefore do not establish a mechanism underlying the prognostic association observed in SHH α tumors, but demonstrate that this association occurs in the context of a gene normally expressed in the relevant cerebellar developmental populations.

Our analysis across different pediatric brain tumor types provides additional context for the association of *PSEN1* with MB biology. *PSEN1* was broadly expressed across the seven tumor entities examined, although expression levels differed significantly among tumor types. MB occupied an intermediate position, with lower *PSEN1* expression than high-grade glioma/astrocytoma, low-grade glioma/astrocytoma, and pilocytic astrocytoma, higher expression than EPN, and levels comparable to those observed in ATRT and diffuse midline glioma. Thus, the favorable prognostic association of higher *PSEN1* expression observed in SHH α MB does not appear to reflect unusually high or MB-specific expression of the gene. Instead, these findings reinforce the notion that the biological significance of *PSEN1* expression is likely to be context dependent. Together with the developmental cerebellar analyses showing *PSEN1* expression across granule cell lineage states, the cross-tumor comparison supports a model in which the prognostic significance of *PSEN1* in SHH α MB may relate more closely to the cellular and transcriptional state in which it is expressed than to its absolute expression level alone.

Several limitations of this study must be acknowledged. First, *PSEN1* transcription levels do not necessarily reflect protein abundance, γ-secretase activity, or functional PS1 signaling. Second, a possible biological significance of *PSEN1* expression in SHH α MB was inferred from transcriptomic and bioinformatic analyses rather than direct experimental measurements, and therefore the present study is purely correlational and cannot establish causality. Likewise, the co-expression networks and enriched biological processes identified here should be considered hypothesis-generating rather than evidence of pathway activation or functional involvement. Currently available MB cell lines do not faithfully reproduce the SHH α molecular context underlying the principal findings reported here, and experiments using appropriately characterized patient-derived or other SHH α-representative models will therefore be required to establish whether *PSEN1* plays a causal role in this MB subtype. Additional studies integrating single-cell, proteomic, functional, and mechanistic approaches will be necessary to determine whether *PSEN1* actively contributes to SHH α MB biology or primarily serves as a marker of underlying cellular states.

The potential value of *PSEN1* as a candidate prognostic marker should be considered in the context of the molecular heterogeneity of MB. Biomarkers that provide information across multiple MB groups, such as *SLFN11*, may offer broader clinical applicability (Nakata et al. 2023). In contrast, the association observed here appears to be restricted to the SHH α molecular context. However, subtype restriction does not necessarily preclude biological or potential clinical relevance, particularly in a disease that is increasingly classified and treated according to molecularly defined entities. Rather, *PSEN1* may represent a candidate marker of biological heterogeneity within SHH α MB. Given the limited number of SHH α cases available for survival analysis, however, the present findings should be considered exploratory and will require confirmation in larger independent cohorts before any clinical application can be proposed.

## 5 Conclusion

The present study uses bioinformatic approaches to identify a possible prognostic association for *PSEN1* in SHH α MB, with elevated expression associated with improved patient survival. Importantly, this association was restricted to a specific molecular subtype, reinforcing the concept that biologically meaningful prognostic factors in MB may be obscured when analyses are performed exclusively at the subgroup level. The *PSEN1*-associated transcriptional pattern identified in SHH α MB was reproduced in an independent cohort, providing additional support for the robustness of this molecular association. Moreover, *PSEN1* was expressed across cell populations of the normal developing human cerebellum, including the granule cell lineage, and across diverse pediatric brain tumor types, with MB displaying intermediate expression levels. Together, these findings suggest that the association of *PSEN1* with favorable outcome in SHH α MB is unlikely to reflect simply selective or unusually high expression of *PSEN1* in this tumor context.

The *PSEN1* expression was also associated with a transcriptional program enriched for intracellular membrane trafficking, endoplasmic reticulum and Golgi function, protein quality control, RNA homeostasis, and developmental signaling pathways including TGF-β and WNT. Combined with the known roles of *PSEN1* in neural progenitor biology and Notch signaling, the developmental cerebellar findings and independent transcriptional validation raise the possibility that *PSEN1* marks a developmentally constrained tumor state characterized by preserved cellular homeostasis and reduced malignant plasticity. Although the mechanistic basis of this association remains to be established, our findings identify *PSEN1* and some of its associated molecular networks as candidates for future functional investigation. More broadly, this study highlights how subtype-specific integrated with developmental and cross-tumor transcriptomic contexts can uncover unexpected biological relationships and generate new hypotheses regarding the developmental programs that influence MB progression and patient outcome.

## Supplementary Information

**Supplementary Table S1.** Analyses of association between high or low *PSEN1* gene expression levels and OS in patients with different subtypes of MB tumors using different statistical criteria.

**Supplementary Table S2.** Comparisons of *PSEN1* expression levels among different molecular subgroups of MB.

**Supplementary Table S3.** Comparisons of *PSEN1* expression levels among different molecular subtypes of MB.

**Supplementary Table S4.** Genes showing strong or very strong co-expression with *PSEN1* in SHH α MB tumors.

**Supplementary Figure S1.** *PSEN1* expression visualized according to age group, survival status, histological subtype, and metastatic status in SHH α MB tumors. The analysis was performed using the R2 Genomics Analysis and Visualization Platform (https://hgserver1.amc.nl/). Data were obtained from the Cavalli cohort (Cavalli et al. 2017); *n* = 65.

**Supplementary Figure S2.** *PSEN1* expression across SHH MB molecular subtypes in an independent cohort. *PSEN1* expression was examined in an independent cohort of molecularly characterized SHH MB using the harmonized expression dataset reported by Arora et al. (2026) and molecular subtype annotations from Skowron et al. (2021). The analysis included 50 SHH α, 42 SHH β, 32 SHH γ, and 72 SHH δ tumors. Differences among subtypes were assessed using the Kruskal–Wallis test (*p* = 0.041), followed by pairwise Wilcoxon rank-sum tests with BH correction for multiple comparisons. Only the difference between SHH β and SHH γ remained significant after BH correction (adjusted *p* = 0.026).

**Supplementary Figure S3.** Independent validation of the *PSEN1*-associated transcriptional pattern in SHH α MB. Pearson correlations between *PSEN1* expression and the prespecified Cavalli-derived *PSEN1*-associated gene set were examined in an independent cohort of 50 SHH α tumors. Of the 26 genes identified as strongly positively correlated with *PSEN1* in the Cavalli cohort, 22 were represented in the independent expression matrix. All 22 genes showed positive correlations with *PSEN1*, with Pearson correlation coefficients ranging from 0.674 to 0.944 and a median correlation coefficient of 0.880.

**Supplementary Figure S4.** Independent validation of *PSEN1* expression across developing human cerebellar RL/EGL populations. *PSEN1* expression was examined in the independent developing human cerebellum single-nucleus RNA-sequencing dataset of Sepp et al. (2024). Cells assigned by the authors to the RL/EGL lineage were grouped according to the original developmental-state annotations. Dot position indicates mean *PSEN1* expression and dot size indicates the percentage of nuclei with detectable *PSEN1* expression. *PSEN1* was detected across granule cell precursor (GCP) and multiple differentiating granule cell (GC) states, independently confirming its expression throughout the developing granule-cell lineage. UBC, unipolar brush cell; UBCP, unipolar brush cell progenitor.

## Author Contributions

**Julia Vanini:** conceptualization, methodology, writing – original draft preparation, data analysis, writing – reviewing and editing. **Amanda Thomaz:** methodology, data analysis, investigation, writing – reviewing and editing. **Marina Müller Lupatini:** conceptualization, methodology, writing – reviewing and editing. **André T. Brunetto:** writing – original draft preparation, writing – reviewing and editing. **Caroline Brunetto de Farias:** methodology, writing – original draft preparation, writing – reviewing and editing. **Mariane Jaeger:** writing – original draft preparation, writing – reviewing and editing. **Rafael Roesler:** conceptualization, methodology, writing – original draft preparation, data analysis, investigation, supervision, writing – reviewing and editing.

## Ethics Statement

This study was based exclusively on analyses of publicly available, de-identified transcriptomic datasets and publicly accessible online bioinformatic resources. No human participants were recruited, no human tissue or biological specimens were collected or analyzed, no identifiable patient information was accessed, and no animal experiments were performed. Therefore, institutional ethics committee approval and informed consent were not required.

## Data Availability Statement

The main tumor dataset used in this study is available in the Gene Expression Omnibus (GEO) repository, accession numbers GSE202043, https://www.ncbi.nlm.nih.gov/geo/query/acc.cgi?acc=GSE202043, GSE85217, https://www.ncbi.nlm.nih.gov/geo/query/acc.cgi?acc=GSE85217, and GSE156053, https://www.ncbi.nlm.nih.gov/geo/query/acc.cgi?acc=GSE156053. The SHH MB RNA-sequencing dataset described by Skowron et al. (2021) is available from the European Genome-Phenome Archive (EGA) under accession number EGAD00001006305 (https://ega-archive.org/datasets/EGAD00001006305). The integrated MB transcriptomic dataset described by Arora et al. (2026) was generated from publicly available RNA-sequencing datasets deposited under accession numbers E-MTAB-6814 (https://www.ebi.ac.uk/biostudies/arrayexpress/studies/E-MTAB-6814), GSE109381 (https://www.ncbi.nlm.nih.gov/geo/query/acc.cgi?acc=GSE109381), EGAS00001002696 (https://ega-archive.org/studies/EGAS00001002696), EGAS00001000254 (https://ega-archive.org/studies/EGAS00001000254), EGAD00001006305 (https://ega-archive.org/dacs/EGAC00001001002), and EGAS00001005826 (https://ega-archive.org/studies/EGAS00001005826). The MB scRNA-seq single cell dataset is available at the Pediatric Neuro-oncology Cell Atlas (https://www.pneuroonccellatlas.org/). Single-nucleus RNA-sequencing data from the developing human cerebellum reported by Aldinger et al. (2021) were obtained through the CELLxGENE collection *Spatial and cell type transcriptional landscape of human cerebellar development* (https://cellxgene.cziscience.com/collections/1b014f39-f202-45ae-bb7d-9286bddd8d8b). The independent developing human cerebellum dataset reported by Sepp et al. (2024) was obtained from the human snRNA-seq dataset in the CELLxGENE collection *Cellular development and evolution of the mammalian cerebellum* (https://cellxgene.cziscience.com/e/bab7432a-5cfe-45ea-928c-422d03c45cdd.cxg/). *PSEN1* expression across pediatric brain tumor types was examined using RNA-sequencing data from the Open Pediatric Brain Tumor Atlas (OpenPBTA; https://github.com/AlexsLemonade/OpenPBTA-analysis).

## Abbreviations

Aβ: Amyloid-β
AD: Alzheimer’s disease
APP: Amyloid precursor protein
ATRT: Atypical teratoid/rhabdoid tumor
BH: Benjamini–Hochberg
BP: Biological Process
CB: Cerebellar tissue
CC: Cellular Component
EGA: European Genome-Phenome Archive
EGL: External granule layer
EPN: Ependymoma
ER: Endoplasmic reticulum
ERAD: ER-associated degradation
FDR: False discovery rate
GCP: Granule cell precursor
GEO: Gene Expression Omnibus
GN: Granule neuron
GO: Gene Ontology
HSD: Honest Significant Difference
iPSC: Induced pluripotent stem cells
MB: Medulloblastoma
MF: Molecular Function
OS: Overall survival
OpenPBTA: Open Pediatric Brain Tumor Atlas
PS1: Presenilin-1
RL: Rhombic lip
scRNA-seq: Single-cell RNA sequencing
SHH: Sonic Hedgehog
SNF: Similarity network fusion
TGF-β: Transforming growth factor-β
TPM: Transcript-per-million
UMAP: Uniform Manifold Approximation and Projection
uRL: Upper rhombic lip
WNT: Wingless

