## Supplementary Information for "*PSEN1* Expression Identifies a Developmentally Distinct Favorable-Prognosis State in SHH α Medulloblastoma"

**SUPPLEMENTARY TABLE S1** | Analyses of association between high or low *PSEN1* gene expression levels and OS in patients with different subtypes of MB tumors using different statistical criteria.

| **Subtype** | **Raw *p* median cutoff** | **Bonferroni *p* scan cutoff** | **Raw *p* scan cutoff** | **Adjusted *p* FDR BH scan cutoff** |
|---|---|---|---|---|
| SHH α | 0.017 | 0.00846 | 2.35e-04 | 0.00282 |
| SHH β | 0.475 | 0.466 | 0.036 | 0.0864 |
| SHH γ | 0.241 | 1.000 | 0.072 | 0.1080 |
| SHH δ | 0.123 | 0.204 | 4.85e-03 | 0.0194 |
| WNT α | 0.330 | 1.000 | 0.231 | 0.2772 |
| WNT β | 0.206 | 0.099 | 0.033 | 0.0864 |
| Group 3 α | 0.666 | 1.000 | 0.221 | 0.2772 |
| Group 3 β | 0.114 | 0.050 | 4.13e-03 | 0.0194 |
| Group 3 γ | 0.906 | 1.000 | 0.365 | 0.3650 |
| Group 4 α | 0.564 | 1.000 | 0.062 | 0.1063 |
| Group 4 β | 0.776 | 1.000 | 0.018 | 0.0720 |
| Group 4 γ | 0.108 | 1.000 | 0.071 | 0.1080 |

FDR, false discovery rate; BH, Benjamini-Hochberg. Data were obtained from the Cavalli cohort (Cavalli et al. 2017).

**SUPPLEMENTARY TABLE S2** ǀ Comparisons of *PSEN1* expression levels among different molecular subgroups of MB.

| | **Diff.** | **Lower** | **Upper** | **Adj. *p*** | **Significance** |
|---|---|---|---|---|---|
| Group 3-SHH | 0.06 | -0.01 | 0.13 | 0.09 | ns |
| Group 4-SHH | 0.04 | -0.02 | 0.09 | 0.36 | ns |
| Group 4-Group 3 | -0.03 | -0.09 | 0.04 | 0.69 | ns |
| Group 3-WNT | -0.08 | -0.17 | 0.02 | 0.15 | ns |
| Group 4-WNT | -0.11 | -0.19 | -0.02 | 0.01 | ** |
| SHH-WNT | -0.14 | -0.23 | -0.05 | 0.00 | *** |

The table shows the difference between pairs, the 95% confidence interval and the *p*-values of the pairwise comparisons. Analyses were performed with Tukey's Honest Significant Difference (HSD) tests. Diff., difference; Adj. *p*, adjusted *p* value. Data were obtained from the Cavalli cohort (Cavalli et al. 2017); ***$p<0.001$; **$p<0.01$; ns, not significant.

**SUPPLEMENTARY TABLE S3** | Comparisons of *PSEN1* expression levels among different molecular subtypes of MB.

| | Diff. | Lower | Upper | Adj. *p* | Significance |
|---|---|---|---|---|---|
| Group 3 α -SHH_γ | 0.28 | 0.13 | 0.43 | 0.00 | *** |
| Group 4 β -Group3 γ | 0.21 | 0.06 | 0.36 | 0.00 | *** |
| Group 3 α -SHH α | 0.21 | 0.07 | 0.35 | 0.00 | *** |
| Group 4 β -SHH γ | 0.17 | 0.03 | 0.32 | 0.00 | ** |
| Group 3 α -SHH δ | 0.16 | 0.02 | 0.29 | 0.01 | ** |
| Group 4 α -Group 3 γ | 0.16 | 0.00 | 0.31 | 0.04 | * |
| Group 3 β-SHH γ | 0.13 | -0.05 | 0.31 | 0.39 | ns |
| Group 4 γ-Group 3 γ | 0.12 | -0.02 | 0.27 | 0.21 | ns |
| SHH δ-SHH γ | 0.12 | -0.03 | 0.27 | 0.24 | ns |
| Group 4 α -SHH γ | 0.12 | -0.02 | 0.26 | 0.22 | ns |
| Group 3 α -SHH β | 0.11 | -0.06 | 0.28 | 0.59 | ns |
| Group 4 β-SHH α | 0.10 | -0.02 | 0.23 | 0.26 | ns |
| SHH β-SHH α | 0.10 | -0.07 | 0.27 | 0.78 | ns |
| Group 4 γ-SHH γ | 0.09 | -0.05 | 0.23 | 0.66 | ns |
| Group 3 α -WNT β | 0.07 | -0.13 | 0.27 | 0.99 | ns |
| Group 3 β-SHH α | 0.06 | -0.11 | 0.23 | 0.99 | ns |
| Group 4 β-Group 4 α | 0.06 | -0.06 | 0.17 | 0.90 | ns |

| | Diff. | Lower | Upper | Adj. *p* | Significance |
|---|---|---|---|---|---|
| Group 4 β-SHH δ | 0.05 | -0.07 | 0.17 | 0.96 | ns |
| SHH δ-SHH α | 0.05 | -0.09 | 0.19 | 0.99 | ns |
| Group 4 α -SHH α | 0.05 | -0.08 | 0.18 | 0.99 | ns |
| Group 4 β-Group 3 β | 0.04 | -0.11 | 0.20 | 1.00 | ns |
| Group 3 α -WNT α | 0.04 | -0.11 | 0.19 | 1.00 | ns |
| Group 4 γ-SHH α | 0.01 | -0.11 | 0.14 | 1.00 | ns |
| Group 3 β-SHH δ | 0.01 | -0.15 | 0.17 | 1.00 | ns |
| Group 4 β-SHH β | 0.01 | -0.15 | 0.16 | 1.00 | ns |
| Group 4 α -SHH δ | -0.00 | -0.13 | 0.12 | 1.00 | ns |
| Group 4 α -Group 3 β | -0.01 | -0.17 | 0.14 | 1.00 | ns |
| WNT β-WNT α | -0.03 | -0.24 | 0.18 | 1.00 | ns |
| Group 4 γ-Group 4 α | -0.03 | -0.14 | 0.08 | 1.00 | ns |
| Group 4 γ-SHH δ | -0.04 | -0.15 | 0.08 | 1.00 | ns |
| Group 4 β-WNT β | -0.04 | -0.23 | 0.16 | 1.00 | ns |
| Group 3 γ-SHH γ | -0.04 | -0.21 | 0.14 | 1.00 | ns |
| Group 3 β-SHH β | -0.04 | -0.23 | 0.15 | 1.00 | ns |
| SHH β-WNT β | -0.04 | -0.26 | 0.18 | 1.00 | ns |

| | Diff. | Lower | Upper | Adj. *p* | Significance |
|---|---|---|---|---|---|
| Group 4 γ-Group 3 β | -0.04 | -0.20 | 0.11 | 1.00 | ns |
| SHH δ-SHH β | -0.05 | -0.21 | 0.12 | 1.00 | ns |
| Group 4 α -SHH β | -0.05 | -0.21 | 0.11 | 1.00 | ns |
| Group 4 β-WNT α | -0.07 | -0.21 | 0.07 | 0.92 | ns |
| SHH β-WNT α | -0.07 | -0.25 | 0.11 | 0.98 | ns |
| SHH γ-SHH α | -0.07 | -0.23 | 0.08 | 0.93 | ns |
| Group 3 β-WNT β | -0.08 | -0.30 | 0.14 | 0.99 | ns |
| Group 4 γ-SHH β | -0.08 | -0.24 | 0.07 | 0.85 | ns |
| Group 4 γ-Group 4 β | -0.09 | -0.19 | 0.02 | 0.24 | ns |
| SHH δ-WNT β | -0.09 | -0.29 | 0.11 | 0.95 | ns |
| Group 4 α -WNT β | -0.09 | -0.29 | 0.10 | 0.93 | ns |
| Group 4 β-Group 3 α | -0.11 | -0.23 | 0.02 | 0.21 | ns |
| Group 3 γ-SHH α | -0.11 | -0.27 | 0.05 | 0.55 | ns |
| Group 3 β-WNT α | -0.11 | -0.29 | 0.07 | 0.66 | ns |
| SHH δ-WNT α | -0.12 | -0.27 | 0.03 | 0.27 | ns |
| Group 4 α -WNT α | -0.12 | -0.26 | 0.02 | 0.17 | ns |
| Group 4 γ-WNT β | -0.12 | -0.32 | 0.07 | 0.61 | ns |

| | Diff. | Lower | Upper | Adj. *p* | Significance |
|---|---|---|---|---|---|
| SHH α -WNT β | -0.14 | -0.34 | 0.06 | 0.53 | ns |
| Group 3 β-Group 3 α | -0.15 | -0.31 | 0.02 | 0.13 | ns |
| Group 4 γ-WNT α | -0.15 | -0.29 | -0.02 | 0.01 | * |
| Group 3 γ-SHH δ | -0.16 | -0.32 | -0.00 | 0.05 | * |
| Group 4 α -Group 3 α | -0.16 | -0.29 | -0.03 | 0.00 | ** |
| Group 3 γ-Group 3 β | -0.17 | -0.35 | 0.02 | 0.12 | ns |
| SHH α -WNT α | -0.17 | -0.32 | -0.02 | 0.02 | * |
| SHH γ-SHH β | -0.17 | -0.35 | 0.01 | 0.09 | ns |
| Group 4 γ-Group 3 α | -0.19 | -0.32 | -0.07 | 0.00 | *** |
| Group 3 γ-SHH β | -0.21 | -0.39 | -0.02 | 0.02 | * |
| SHH γ-WNT β | -0.21 | -0.42 | 0.00 | 0.05 | ns |
| SHH γ-WNT α | -0.24 | -0.41 | -0.08 | 0.00 | *** |
| Group 3 γ-WNT β | -0.25 | -0.47 | -0.03 | 0.01 | * |
| Group 3 γ-WNT α | -0.28 | -0.45 | -0.11 | 0.00 | *** |
| Group 3 γ-Group 3 α | -0.32 | -0.48 | -0.15 | 0.00 | *** |

The table shows the difference between pairs, the 95% confidence interval and the *p*-values of the pairwise comparisons. Analyses were performed with Tukey's Honest Significant Difference (HSD) tests. Diff., difference; Adj. *p*, adjusted *p* value. Data were obtained from the Cavalli cohort (Cavalli et al. 2017); ***$p<0.001$; **$p<0.01$; *$p<0.05$; ns, not significant.

**SUPPLEMENTARY TABLE S4** | Genes showing strong or very strong co-expression with *PSEN1* in SHH α MB tumors.

| **Rank** | **Gene** | **Spearman *r*** |
|---|---|---|
| 1 | *SPTLC2* | 0.7376 |
| 2 | *ATL1* | 0.7365 |
| 3 | *EXD2* | 0.7353 |
| 4 | *ZNF410* | 0.7300 |
| 5 | *PAPOLA* | 0.7014 |
| 6 | *SOS2* | 0.6947 |
| 7 | *EXOC5* | 0.6867 |
| 8 | *VIPAR* | 0.6662 |
| 9 | *SEL1L* | 0.6651 |
| 10 | *SLC39A9* | 0.6611 |
| 11 | *MUDENG* | 0.6570 |
| 12 | *C14ORF118* | 0.6513 |
| 13 | *TMED8* | 0.6449 |
| 14 | *SETD3* | 0.6447 |
| 15 | *WDR20* | 0.6442 |
| 16 | *C14ORF101* | 0.6427 |
| 17 | *SNX6* | 0.6421 |
| 18 | *DCAF5* | 0.6381 |
| 19 | *GOLGA5* | 0.6379 |
| 20 | *PPM1A* | 0.6365 |
| 21 | *ACTR10* | 0.6347 |
| 22 | *TRAPPC6B* | 0.6303 |
| 23 | *SNW1* | 0.6276 |
| 24 | *ZFYVE1* | 0.6271 |
| 25 | *ZC3H14* | 0.6251 |
| 26 | *JKAMP* | 0.6210 |

Data were obtained from the Cavalli cohort (Cavalli et al. 2017); $n = 65$; all $p$s<0.05.

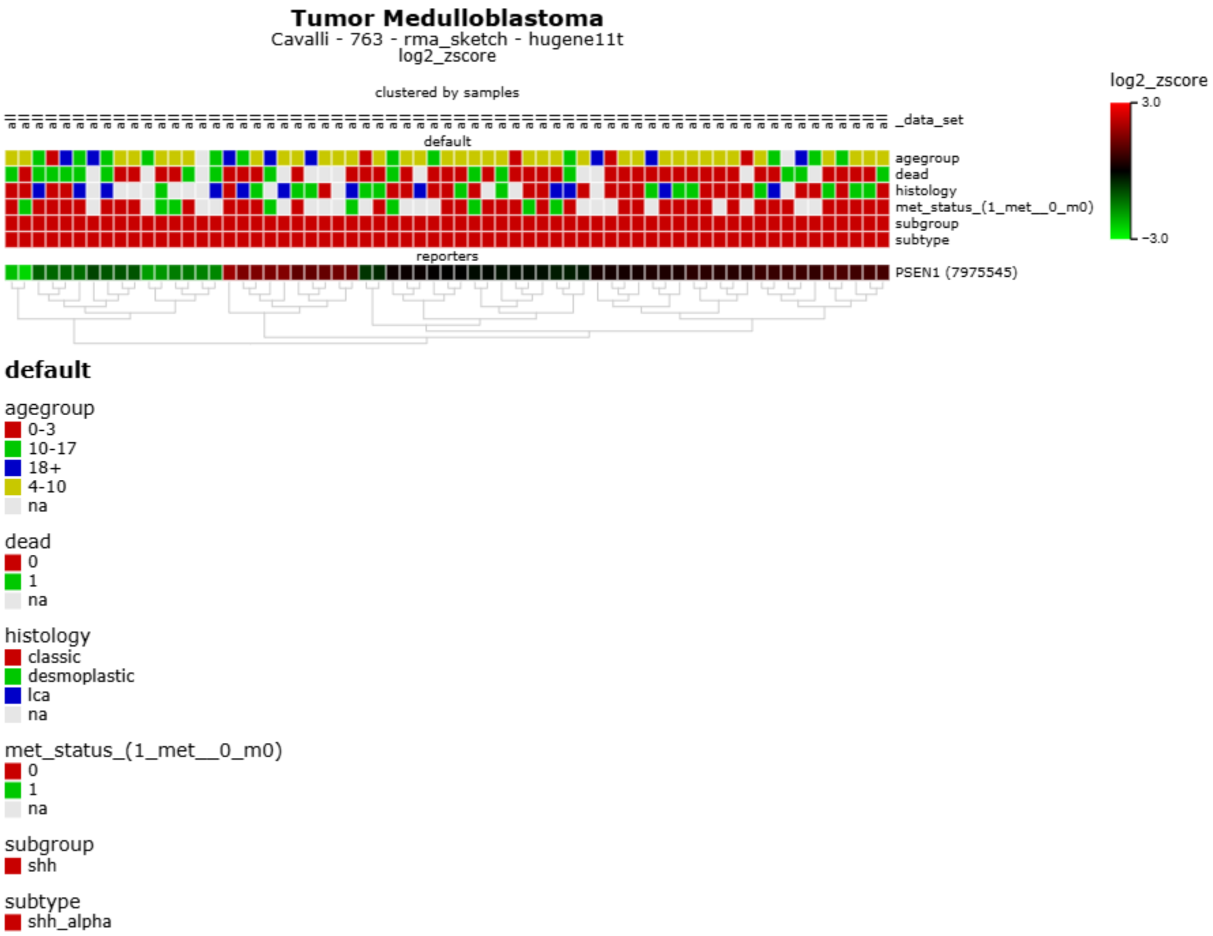


**SUPPLEMENTARY FIGURE S1** | *PSEN1* expression visualized according to age group, survival status, histological subtype, and metastatic status in SHH α MB tumors. The analysis was performed using the R2 Genomics Analysis and Visualization Platform (https://hgserver1.amc.nl/). Data were obtained from the Cavalli cohort (Cavalli et al. 2017); *n* = 65.

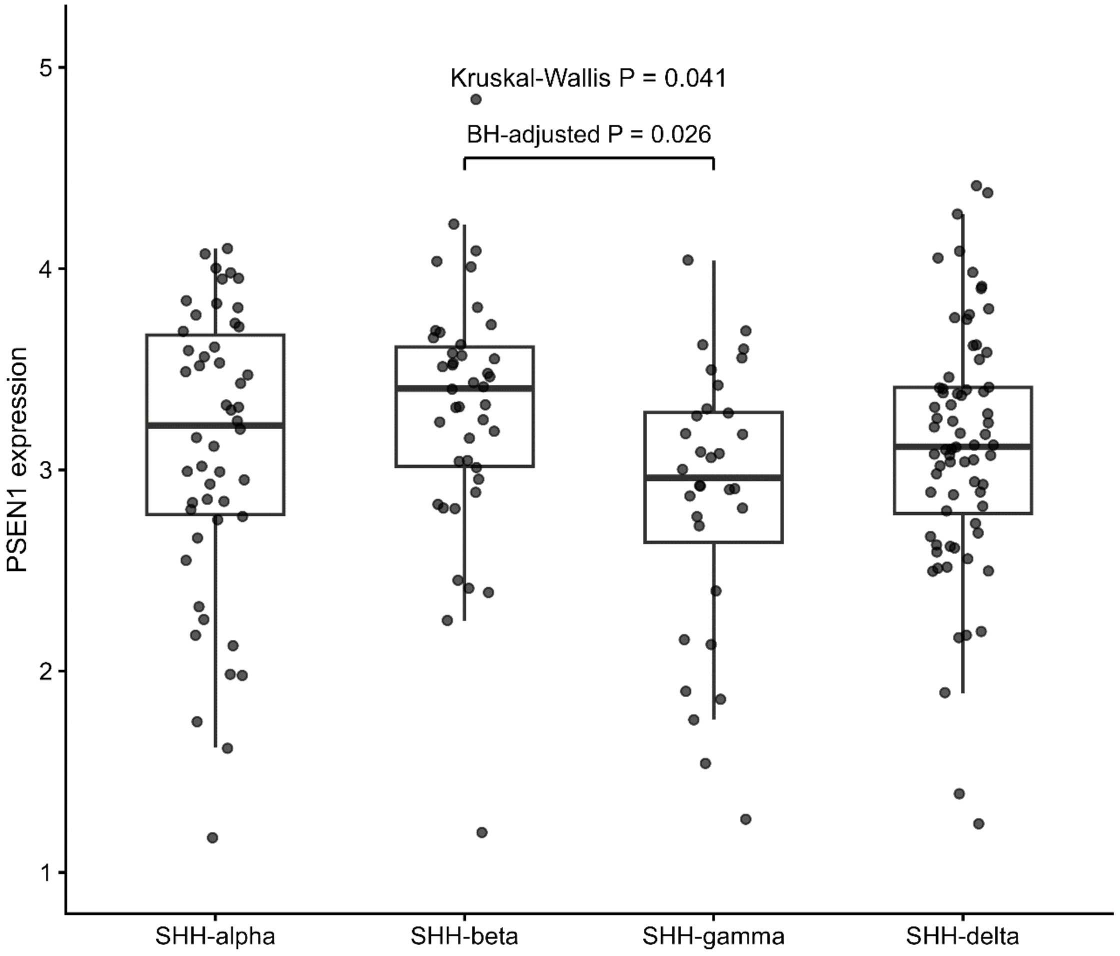


**SUPPLEMENTARY FIGURE S2** I *PSEN1* expression across SHH MB molecular subtypes in an independent cohort. *PSEN1* expression was examined in an independent cohort of molecularly characterized SHH MB using the harmonized expression dataset reported by Arora et al. (2026) and molecular subtype annotations from Skowron et al. (2021). The analysis included 50 SHH α, 42 SHH β, 32 SHH γ, and 72 SHH δ tumors. Differences among subtypes were assessed using the Kruskal–Wallis test ($p$ = 0.041), followed by pairwise Wilcoxon rank-sum tests with BH correction for multiple comparisons. Only the difference between SHH β and SHH γ remained significant after BH correction (adjusted $p$ = 0.026).

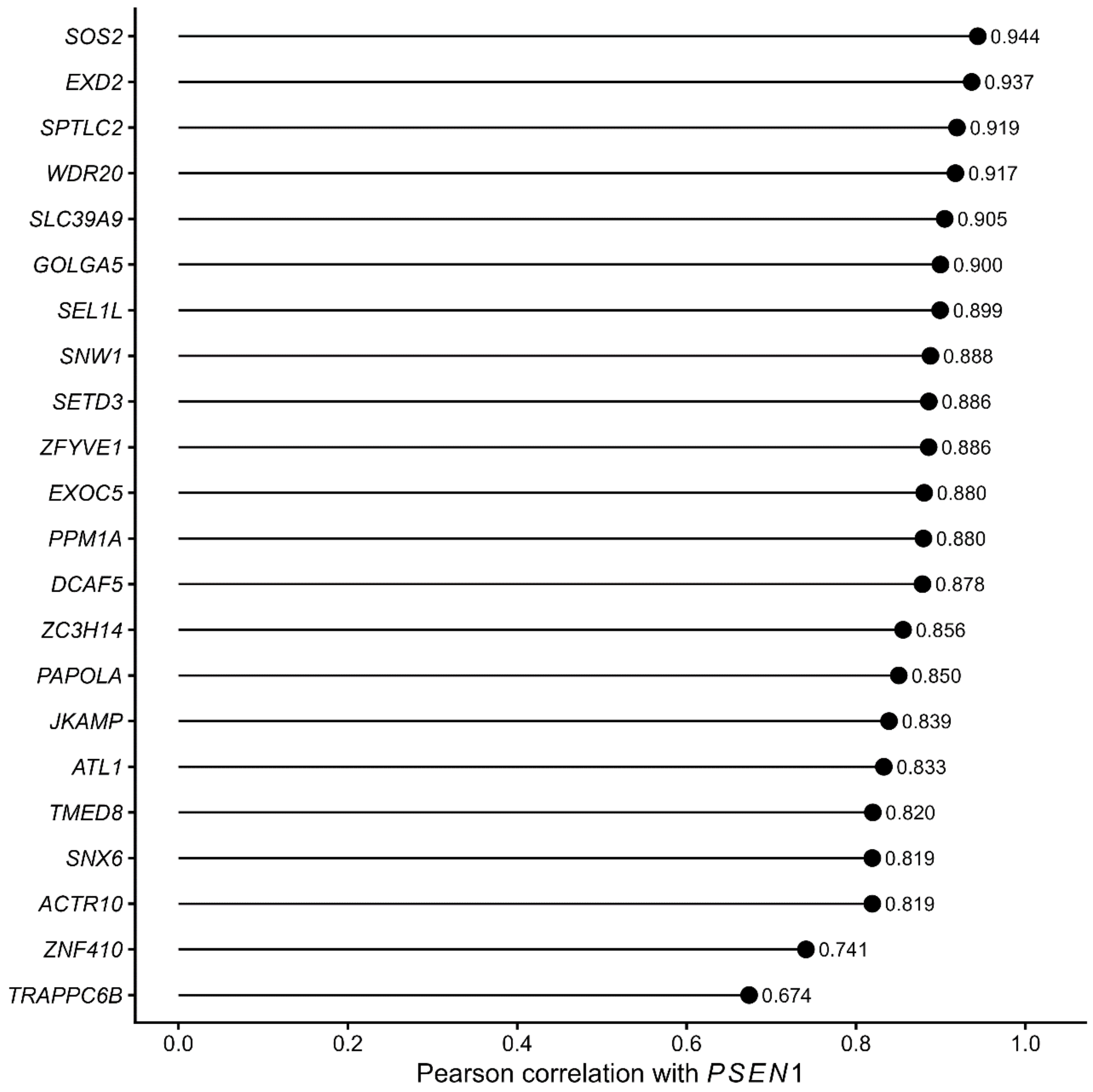


**SUPPLEMENTARY FIGURE S3** | Independent validation of the *PSEN1*-associated transcriptional pattern in SHH α MB. Pearson correlations between *PSEN1* expression and the prespecified Cavalli-derived *PSEN1*-associated gene set were examined in an independent cohort of 50 SHH α tumors. Of the 26 genes identified as strongly positively correlated with *PSEN1* in the Cavalli cohort, 22 were represented in the independent expression matrix. All 22 genes showed positive correlations with *PSEN1*, with Pearson correlation coefficients ranging from 0.674 to 0.944 and a median correlation coefficient of 0.880.

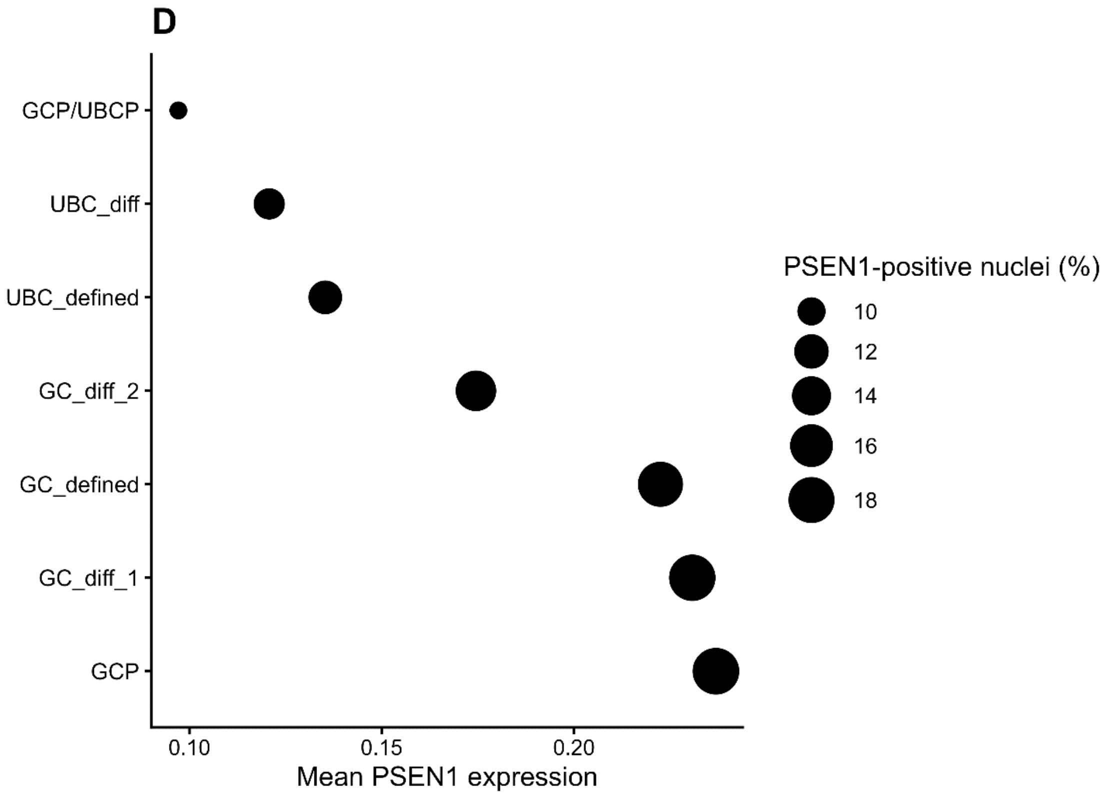


**SUPPLEMENTARY FIGURE S4** | Independent validation of *PSEN1* expression across developing human cerebellar RL/EGL populations. *PSEN1* expression was examined in the independent developing human cerebellum single-nucleus RNA-sequencing dataset of Sepp et al. (2024). Cells assigned by the authors to the RL/EGL lineage were grouped according to the original developmental-state annotations. Dot position indicates mean *PSEN1* expression and dot size indicates the percentage of nuclei with detectable *PSEN1* expression. *PSEN1* was detected across granule cell precursor (GCP) and multiple differentiating granule cell (GC) states, independently confirming its expression throughout the developing granule-cell lineage. UBC, unipolar brush cell; UBCP, unipolar brush cell progenitor.
